# Approaching an Error-Free Diploid Human Genome Using a Support-Based Validation Framework

**DOI:** 10.1101/2025.08.01.667781

**Authors:** Yanan Chu, Zhuo Huang, Changjun Shao, Shuming Guo, Yiji Yang, Xinyao Yu, Yurong Luo, Jian Wang, Yabin Tian, Jing Chen, Ran Li, Yukun He, Stylianos E. Antonarakis, Jun Yu, Jie Huang, Zhancheng Gao, Yu Kang

## Abstract

Complete human genomes now resolve regions long inaccessible to genetic analysis, yet repetitive and structurally complex sequences remain prone to assembly errors because of limitations in current assembly algorithms. We introduce a support-based framework founded on the principle that each sequencing read provides independent evidence for the underlying genome sequence. Implemented through Sufficient Alignment Support (SAS), the framework identifies regions lacking concordant read support, localizes residual errors, and guides correction. Applying SAS to T2T-YAO, a haplotype-resolved Han Chinese genome, produced a support-validated assembly in which nearly all sequences outside unresolved rDNA arrays and long homopolymer tracts are supported by independent sequencing evidence. This work establishes a scalable framework for genome validation and provides a support-validated East Asian diploid reference for investigating human genomic diversity.

## Main Text

The advent of telomere-to-telomere (T2T) human genome assemblies has enabled the generation of highly complete diploid genomes across diverse populations (*1-5*). These assemblies extend genomic analysis into previously inaccessible regions, including centromeres (*6*), segmental duplications (*7*), acrocentric short arms and other repetitive sequences that together with other common repeats (*8*) comprise more than half of the human genome. As population-scale pangenome projects expand, these regions are increasingly recognized as contributors to genome evolution, gene regulation, and human disease (*9-12*). They are among the most rapidly evolving components of the human genome and exhibit extensive structural diversity among individuals and populations (*13, 14*).

The growing number of complete human genomes is rapidly expanding the catalog of sequences absent from GRCh38 and other existing references, many of which originate from repetitive and structurally complex regions (*15-17*). Yet these regions remain particularly susceptible to assembly errors because current assembly algorithms still struggle to resolve highly similar repeats, large segmental duplications, and structurally complex loci. As a result, the correctness of many newly assembled sequences cannot be assumed despite the apparent completeness of modern genome assemblies. Consequently, as genome assembly approaches completeness, establishing genome correctness has become the next major challenge for human genomics (*18*).

A central obstacle is the absence of a consistent framework for defining and evaluating genome correctness, particularly in complex genomic regions. Existing assessment methods were largely developed before the advent of T2T genomes, when repetitive regions were incomplete or inaccessible. These approaches typically rely on either k-mer–based completeness estimates (*19*) or read–assembly discrepancy signals. Although effective in unique sequences, both become increasingly difficult to interpret in repetitive regions. K-mer–based metrics may fail to detect assembly errors when structurally distinct repeat copies retain similar k-mer profiles, while discrepancy-based methods are strongly influenced by alignment ambiguity and sequencing biases. As a result, the genomic regions that contribute most to human structural diversity are often the regions for which assembly correctness is most difficult to establish.

We address this challenge by introducing a support-based framework that evaluates genome assemblies using direct experimental evidence. The framework is based on the principle that each sequencing read provides independent support for the underlying biological sequence, and that genomic regions lacking sufficient concordant support across complementary datasets likely reflect potential residual assembly errors. We implement this framework through Sufficient Alignment Support (SAS), which efficiently localizes both structural and nucleotide-level inconsistencies and enables systematic assembly refinement. To improve genome correctness across genomic scales, we applied a hierarchical correction strategy that progresses from megabase-scale structure to kilobase-scale organization and ultimately nucleotide-level sequence accuracy using complementary technologies. Conceptually, this approach parallels the hierarchical clone-by-clone strategy of the Human Genome Project, in which physical maps, BAC-libraries, and Sanger sequencing were integrated to progressively establish genome correctness across increasingly finer scales.

Applying SAS and SAS-guided correction to T2T-YAO v1.1, a haplotype-resolved telomere-to-telomere genome of a Han Chinese individual, we generated T2T-YAO v2.0, a support-validated diploid genome in which nearly all genomic sequences outside unresolved rDNA arrays and a limited number of long homopolymer tracts are supported by independent sequencing evidence. Orthogonal evaluation confirmed improved read-assembly consistency across centromeres, segmental duplications, multicopy genes, and other structurally complex regions. Together, this work establishes a scalable framework for experimentally grounded genome validation and correction. The resulting support-validated genome provides a benchmark resource for evaluating genome analysis in repetitive regions and a population-relevant diploid reference for investigating complex human genomic diversity.

## Results

### Support-based assembly correctness

To systematically evaluate residual inconsistencies in a telomere-to-telomere diploid human genome, we generated a comprehensive multi-platform sequencing dataset from YAO-derived induced pluripotent stem cells (iPSCs), including ultra-long Oxford Nanopore (ONT) R10 reads (*20*), Pore-C (*21*), high-fidelity SPRQ reads and high-accuracy Element Q50 reads (*22*) (table S1). These datasets provide complementary strengths in read length, structural continuity, and base accuracy, enabling systematic assessment of residual assembly errors under current state-of-the-art sequencing platforms.

We first evaluated the previously released T2T-YAO v1.1 assembly (*2*) using existing k-mer– based and discrepancy-based approaches. These methods produced markedly inconsistent estimates of residual errors (Fig. 1A). Merqury-based evaluation (*19, 23*) using a hybrid Element–SPRQ k-mer set, estimated approximately 10^3^–10^4^ residual errors across the diploid genome. In contrast, discrepancy-based methods—including DeepVariant (*24, 25*), GATK HaplotypeCaller (*26*) and DeepPolisher (*27*), Sniffles2 (*28*), Flagger (*15*) and nucflag (*29*)— identified substantially larger numbers of discrepancies between reads and assembly, including 63,398 non-redundant base-level discrepancies, 25.61 Mb of abnormal coverage regions and 1,050 structural disruptions. Notably, most discrepancy signals detected by these methods enriched in repetitive regions, such as centromeres and telomeres. These discrepancies that do not generate novel k-mers remain invisible to k-mer–based evaluation. Conversely, we found that some k-mer–derived error signals were enriched in regions lacking SPRQ and Element coverage, suggesting that they arise from sequencing biases rather than true assembly errors.

**Fig. 1.**
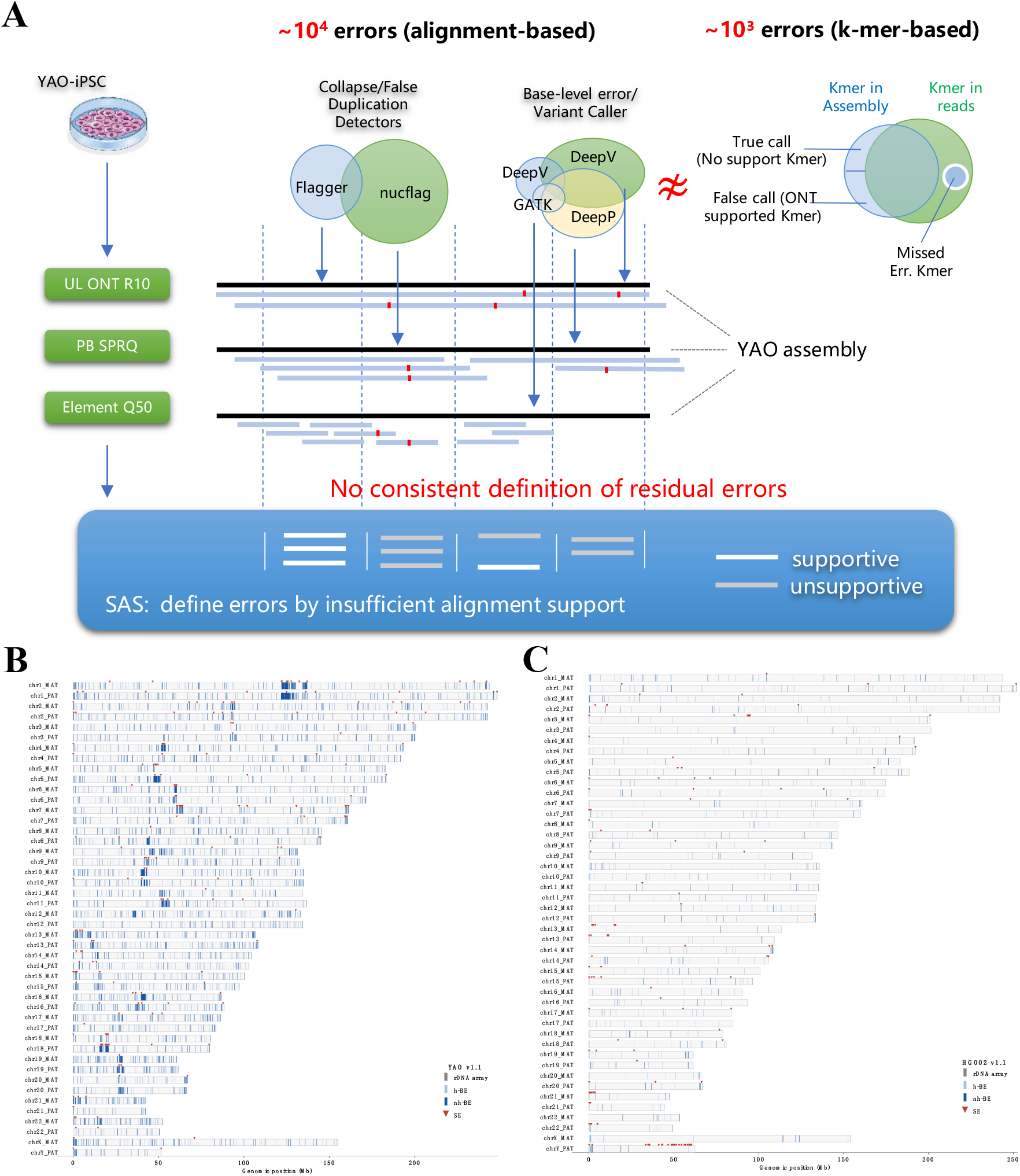
Support-based definition of residual error and assembly correctness. (**A**) Multiple orthogonal sequencing technologies (short reads, HiFi, ONT) and commonly used error-detection methods (SV and SNV callers, k-mer–based evaluation) yield largely discordant error sets when applied to the same assembly, reflecting differences in alignment signals and error definitions across platforms and tools. We introduce a support-based framework for defining assembly correctness, in which correctness is determined by consistent support from independent sequencing reads. Regions lacking sufficient or coherent alignment support are considered candidate errors, providing a unified, evidence-based criterion for genome assembly evaluation.Red dot line read represent mismatch between read and assembly. b, c, Genome-wide application of SAS reveals the distribution of residual inconsistencies in YAO v1.1 (**B**) and the benchmark assembly HG002 v1.1 (**C**), with signals enriched in repetitive regions and substantially reduced in HG002.

Limited concordance was also observed among discrepancy-based detectors themselves. Across base-level callers, only 126 discrepancy signals were shared by all methods despite thousands of candidate calls reported individually. Similarly, coverage-based detectors exhibited substantial inconsistencies in regions that were either erroneously collapsed or duplicated in the assembly (data S2). These observations indicate that existing approaches frequently generate fragmented and partially conflicting residual error signals, particularly within repetitive genomic regions, where platform-specific sequencing characteristics, alignment ambiguity and detector-dependent thresholds contribute to substantial false-positive and false-negative assembly error predictions.

This inconsistency reflects both the absence of a consistent framework for defining “assembly correctness” and limitations of current approaches in accurately localizing residual assembly errors. As each sequencing read represents independent evidence for the presence of genomic sequence within a DNA sample, we reasoned that assembly correctness could be defined through direct sequencing support instead of read-assembly discrepancy. Under this framework, genomic regions lacking sufficient support from concordant sequencing reads are defined as assembly inconsistencies, which may indicate potential residual assembly errors (Fig. 1A). Importantly, this support-based framework relies on experimental evidence than assumptions or machine-learning training to distinguish true assembly errors from sequencing errors, thereby avoiding circular validation logic in which assemblies are evaluated using the same datasets used for assembly construction.

Based on this principle, we developed Sufficient Alignment Support (SAS), a support-based framework that screens genomic region in a window-by-window manner for sufficient supported from aligned reads generated by at least one sequencing platform (fig. S1A). Rather than reconciling heterogeneous discrepancy signals across methods, SAS directly evaluates the presence or absence of local sequencing support. Base-level inconsistencies are detected using 50 bp windows integrating data from all sequencing platforms, whereas structural inconsistencies are assessed using 2 kb windows derived from Oxford Nanopore long reads. Additional operational definitions of alignment support, criteria for length of homopolymer tracts and abnormal depth are described in Methods and fig. S2. In benchmarking analyses using simulated spike-in errors, SAS achieved F1 scores exceeding 0.99 and substantially outperformed existing discrepancy-based approaches (Methods, fig.S1B, data S3).

Application of SAS to T2T-YAO v1.1 identified 21,511 base-level windows and 4,438 structural windows lacking of support. These unsupported regions were enriched within structurally complex genomic regions, including ribosomal DNA (rDNA) arrays and centromeres (Fig. 1B, data S4). Notably, most of the high-confidence discrepancy signals reproducibly detected by existing variant-, structure- and coverage-based methods were contained within SAS-defined unsupported regions, indicating that diverse residual error signals can be unified under a single support-based framework. We also noticed that long homopolymer tracts (PolyA/T ⩾10, PolyC/G ⩾7 and dinucleotide repeats ⩾10) exhibited substantial inconsistencies among reads generated by ONT and SPRQ platforms, suggesting that these regions may remain intrinsically difficult to resolve using current sequencing technologies (fig. S2B). We therefore classified these long homopolymer-associated signals (h-BEs) separately from technically resolvable non-homopolymer-associated inconsistencies (nh-BEs).

We next applied SAS to HG002 v1.1(4) and identified substantially fewer unsupported regions, consistent with its higher baseline quality (Fig. 1C). Together, these results demonstrate that support-based evaluation provides a unified and experimentally grounded framework for identifying residual assembly inconsistencies across both base-level and structural scales.

### YAO v2.0: first support-validated genome

To determine whether support-based evaluation could guide the identification of regions requiring correction and ultimately enable a complete human genome assembly, we started from T2T-YAO v1.1 and implemented an SAS-guided correction framework in which unsupported regions were selectively refined. Because assembly inconsistencies elsewhere can induce read misalignments that obscure the true sequence of some unsupported regions, only a subset of problematic loci can typically be resolved in a single iteration. Repeated cycles of correction, read realignment, and support reassessment progressively exposed additional resolvable loci, enabling convergence toward a support-validated assembly (fig. S3). In contrast, convergence is difficult to achieve with existing evaluation approaches, which often generate spurious signals that hinder the reliable localization of residual assembly errors and therefore impede iterative refinement. Accurate identification of regions that genuinely require correction is a fundamental prerequisite for assembly convergence.

Meanwhile, our correction strategy was inspired by the hierarchical clone-by-clone paradigm of the Human Genome Project, in which physical maps, BAC-libraries, and Sanger sequencing progressively refined genome accuracy from chromosomal structure to nucleotide sequence. Analogously, we combined Pore-C chromatin contact maps, ultra-long Oxford Nanopore reads, and high-accuracy sequencing data to guide correction at megabase-, kilobase-, and nucleotide-scale resolution, respectively (Fig. 2A). By enforcing a correction order in which large structural resolution precedes local refinement, this hierarchical strategy minimizes alignment-induced artifacts and improves discrimination of repetitive and duplicated sequences.

**Fig. 2.**
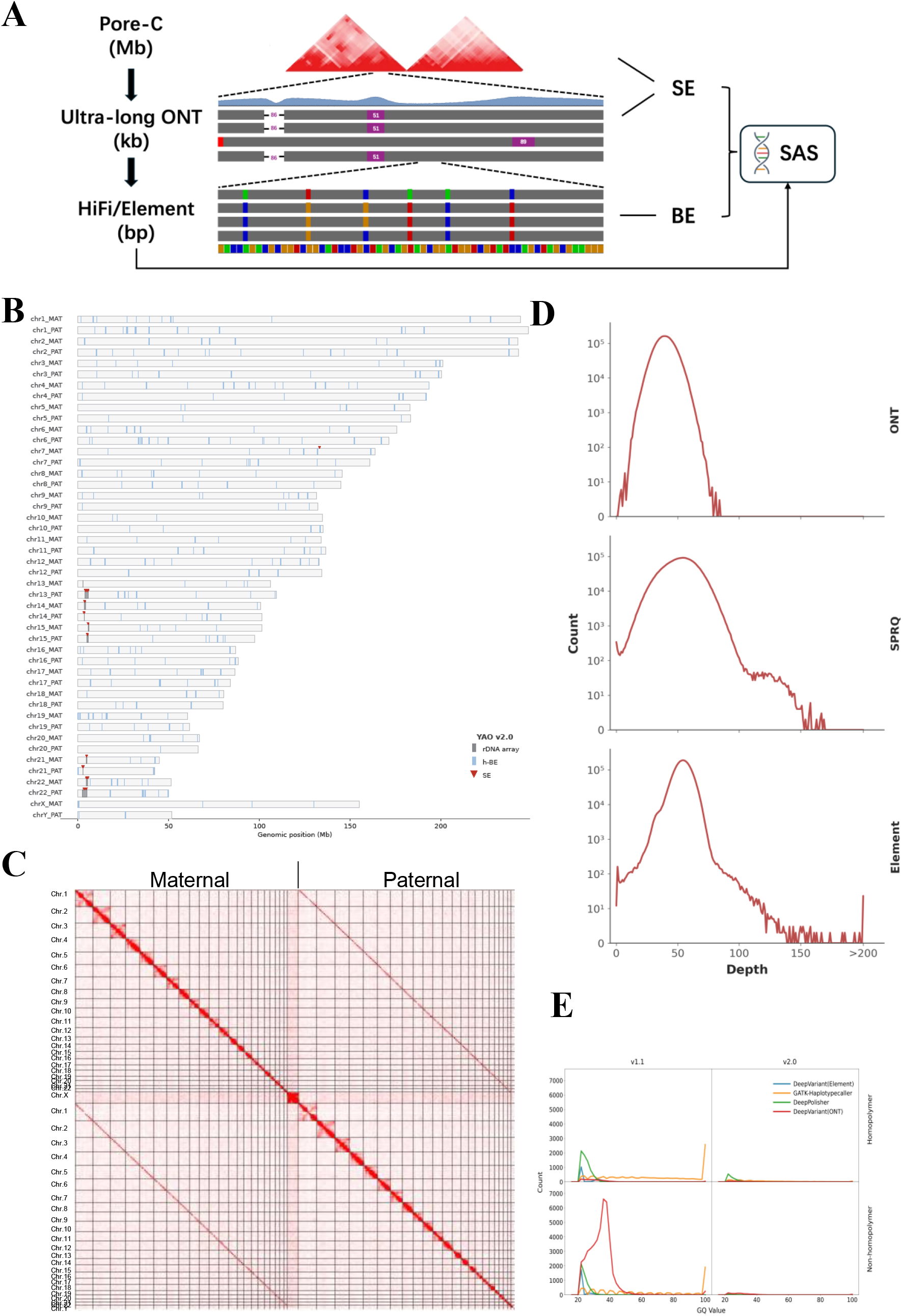
T2T-YAO v2.0, the first support-validated human genome resource. (**A**) Hierarchical genome correction framework integrating scale-resolved sequencing evidence, where structural, continuity, and nucleotide-level errors are resolved sequentially using Pore-C, ultra-long nanopore reads, and high-accuracy sequencing, respectively. SE, structural error; BE, base-level error. (**B**) The final T2T-YAO v2.0 assembly shows complete alignment support across all resolvable genomic regions, with remaining unsupported signals confined to ribosomal DNA arrays and 348 long homopolymer tracts. (**C**) Pore-C contact map of T2T-YAO v2.0 showing long-range chromosomal organization, including clear haplotype separation and correct pairing of acrocentric chromosome arms, supporting accurate structural assembly. (**D**) Distribution of read depth across ONT, SPRQ, and Element sequencing platform in T2T-YAO v1.1 and v2.0 outside rDNA regions. ONT read depth follows a near-normal distribution and provides continuous genome-wide coverage, whereas SPRQ and Element data exhibit skewed coverage distributions and localized dropout regions. (**E**) Distribution of genotype quality (GQ) scores for SNV-like discrepancies identified by multiple variant callers in T2T-YAO v1.1 and v2.0, stratified by homopolymer and non-homopolymer contexts. Discrepancy signals are markedly reduced in v2.0, indicating elimination of spurious variant calls arising from alignment ambiguity.

To minimize over-correction in base-level errors, we developed a platform-integrated window consensus (PWC) framework that restricts sequence refinement to SAS-defined unsupported windows. Candidate consensus sequences were generated independently from multiple sequencing platforms and prioritized according to their platform-specific error profiles. Because h-BEs (base errors within long homopolymer tracts) remain challenging to resolve reliably with current long-read sequencing technologies, corrections were applied only when supported by Element sequencing data. In contrast, nh-BEs were iteratively refined using complementary evidence from all sequencing platforms (Methods, fig. S2B).

Iterative application of SAS-guided correction progressively reduced unsupported regions and converged after 47 rounds of polishing. In the final T2T-YAO v2.0 assembly, nearly all non-rDNA genomic regions satisfied SAS-defined structural support criteria, with a single exception: a 12-kb region showing marginally elevated local depth relative to the predefined SAS threshold (Fig. 2B). However, read alignments at this locus show no evidence of disrupted continuity or chromosomal misassembly, suggesting that this signal likely reflects local mapping ambiguity in highly repetitive sequences rather than true structural inconsistency (fig. S4A). At the base level, all non-homopolymer-associated unsupported regions outside rDNA arrays were resolved in T2T-YAO v2.0. The remaining unsupported loci outside rDNA clusters consisted of 348 long homopolymer-tracts, accounting for only 0.014% of all genome-wide loci (Fig. 2B). These loci were strongly enriched in highly repetitive regions, including the D4Z4 repeat array (*30*), where accurate repeat-length determination remains limited by current sequencing technologies (fig. 4B). Together, these results indicate that SAS-guided refinement enables systematic elimination of resolvable residual inconsistencies, yielding a support-validated human genome under current technological constrains.

To independently evaluate the residual inconsistency level of T2T-YAO v2.0, we reperformed multiple orthogonal assessment methods, including k-mer–based, structure-based and base-level discrepancy analyses, and compared the results with both the original T2T-YAO v1.1 assembly (data S1) and the benchmark assembly HG002 v1.1 (*4*) (Table 1).

**Table 1.**
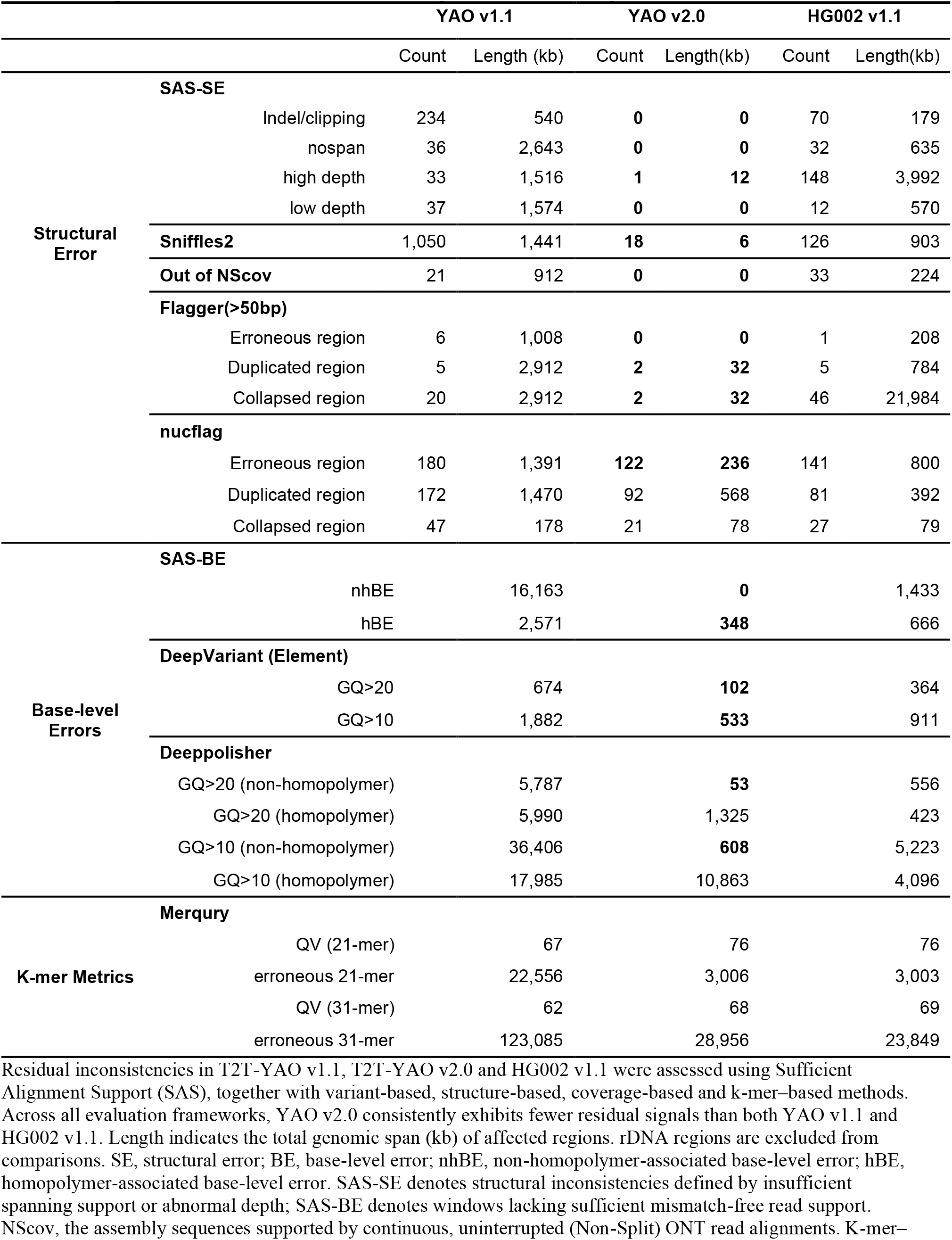
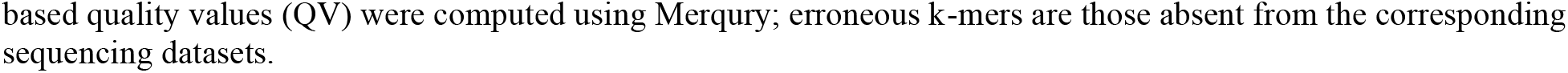
Comparative evaluation of residual error signals across human genome assemblies.

Merqury analysis showed substantial improvement in k-mer–based quality following SAS-guided refinement, with QV increasing from 67 to 76 for 21-mers and from 62 to 68 for 31-mers, almost equal to the reported quality of HG002 v1.1. Residual erroneous k-mers in T2T-YAO v2.0 were highly localized and predominantly enriched within low-complexity or repetitive sequence contexts, including GA/CT-rich tracts, GC-rich tandem repeats and non-canonical telomeric motifs, which are known to exhibit sequencing bias across current technologies (*31, 32*) (data S5). Incorporation of ONT-derived k-mers into a trinary k-mer set eliminated all remaining erroneous k-mers, yielding no detectable erroneous 21-mers (table S2) indicating that residual k-mer signals primarily reflect sequence-intrinsic sequencing limitations rather than assembly errors.

Long-range structural evaluation using Pore-C contact maps confirmed accurate chromosomal organization, haplotype phasing and acrocentric chromosome architecture in T2T-YAO v2.0 (Fig. 2C). Consistently, all non-rDNA genomic regions were covered by uninterrupted non-split ONT read alignments (NScov), supporting genome-wide structural integrity (Fig. 2D, Table 1). Coverage-based analyses using Flagger identified only a small number of abnormal-depth regions, more than 99% of which were confined to rDNA arrays and telomeric regions (fig. S5, data S2). In contrast to the near-normal distribution of ONT coverage, SPRQ and Element exhibit skewed coverage distribution and localized dropout regions (Fig. 2D). Under the same evaluation framework, HG002 v1.1 showed substantially higher numbers and broader spans of residual structural signals than T2T-YAO v2.0 (Table 1).

Base-level evaluation using DeepVariant, DeepPolisher and GATK HaplotypeCaller similarly showed marked reduction of discrepancy signals in T2T-YAO v2.0. The total number of high-confidence SNV-like calls (GQ >20) from all methods decreased from 63,398 in YAO v1.1 to 2,354 in YAO v2.0, (Fig. 2E). Most remaining calls corresponded homopolymer tracts flagged by DeepPolisher, while only 38 were non-homopolymer-associated discrepancies. Of these, 36 were located within rDNA regions, and the remaining two were supported by sufficient read evidence consistent with the assembly sequence upon manual inspection.

Across all orthogonal evaluation frameworks, T2T-YAO v2.0 consistently exhibited the lowest level of residual discrepancy signals among all evaluated assemblies (Table 1). Together, these results establish T2T-YAO v2.0 as a support-validated reliable human genome resource and demonstrate that genome assembly inconsistencies can be systematically resolved using direct experimental support under current sequencing constraints.

### Support-validated repetitive regions in human genome

The high structural continuity and minimal residual inconsistencies of T2T-YAO v2.0 enabled reliable characterization of highly repetitive genomic regions that remain difficult to resolve in conventional human genome assemblies, including centromeres, tandem repeats, multicopy genes and long homopolymer loci.

We first characterized segmental duplication (SD) regions in T2T-YAO v2.0, which span 188.6 Mb and 181.2 Mb in the maternal and paternal haplotype, respectively, substantially less than the 207.6 Mb reported in T2T-CHM13 (*7*). SD regions exhibited strong synteny between the two YAO haplotypes, but apparently more disrupted synteny relative to T2T-CHM13, together with increased SD-nonSD inter-alignment, suggesting extensive structural diversity within duplicated regions across human genomes (Fig. 3A). To assess gene completeness within repetitive and duplicated regions, we transferred gene annotations from T2T-CHM13 v2.0 using Liftoff (*33*). Excluding highly divergent rDNA clusters and immune receptor loci, 56,601 (98.65%) of 57,377 annotated genes were successfully mapped to at least one haplotype of T2T-YAO v2.0 table S3). A total of 112 protein-coding genes remained unmapped in both haplotypes even after additional recovery using protein-to-genome alignment (*34*) against the MANE v1.4 database (*35*), whereas lncRNAs and pseudogenes exhibited even higher loss rates. Annotation of T2T-YAO v2.0 further identified 3,012 and 4,112 additional copidaes of 1,193 non-redundant muti-copy genes in maternal and paternal haplotype, respectively, comparable to the ∼3500 additional genes identified in T2T-CHM13 (*1*). These included protein-coding genes, lncRNAs, and pseudogenes distributed throughout the diploid genome, of which 64.18% resided within SD regions (Fig.3B, data S6). However, 959 of the 1,193 non-redundant muti-copy genes identified in T2T-YAO displayed copy number differences relative to T2T-CHM13, further supporting substantial structural diversity within duplication regions among human genomes. Importantly, although SD regions in T2T-YAO collectively constitute 6.23% of the genome and contain 5,668 genes (∼10% of all annotated human genes), they account for only 2.16% ClinVar records lifted over from GRCh38, highlighting the persistent difficulty of identifying disease-associated variants within duplicated regions because of ambiguous or erroneous read alignments.

**Fig. 3.**
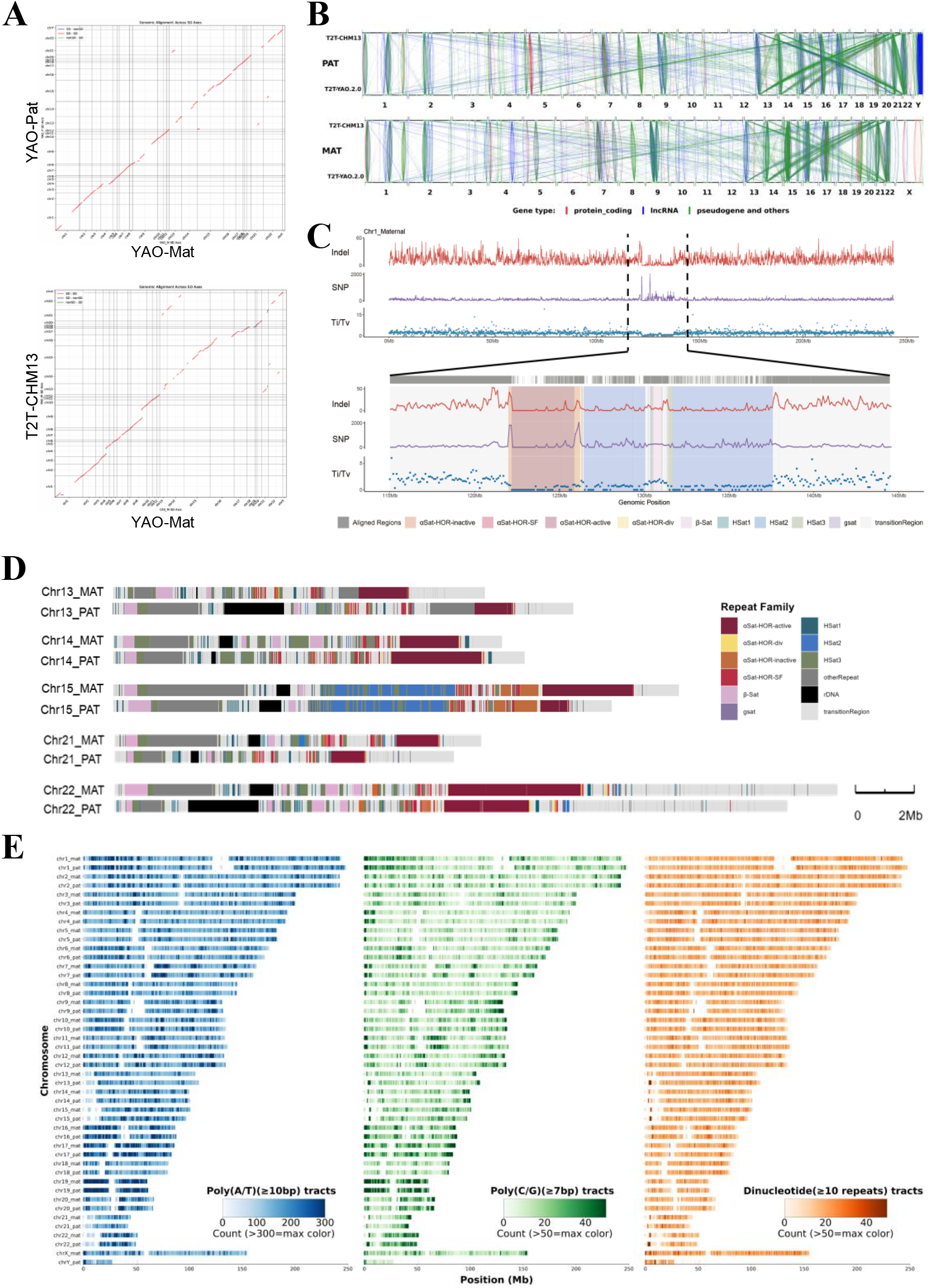
Support-validated repetitive regions in the diploid T2T-YAO v2.0 assembly. (**A**) Dot-plot comparison of segmental duplication (SD) regions between the maternal and paternal haplotypes of T2T-YAO v2.0 (top) and between T2T-YAO maternal haplotype and T2T-CHM13 (bottom). The two YAO haplotypes exhibit substantially higher synteny than comparisons with T2T-CHM13, indicating structural diversity of SD regions among human genomes. (**B**) Genome-wide landscape of potential multi-copy gene expansions in the T2T-YAO v2.0 genome. This connectivity map illustrates the distribution of amplified gene sets relative to the T2T-CHM13 reference and are colored by gene type. High-confidence secondary alignments (sequence identity >0.95) were utilized to identify putative novel gene copies from ‘one-to-many’ redundant mappings and anchored to their original CHM13 loci via Bézier curves. (**C**) Density of SNPs and small indels, and transition/transversion (Ti/Tv) ratio in 100 kb windows along paternal chromosome 1. Upper panel: whole chromosome; lower panel: expanded view of centromere and flanking regions, with repeat annotations indicated in the background of the peak map. (**D**) Repeat organization of the resolved short arms of acrocentric chromosomes in both haplotypes with distinct satellite repeat subfamilies and transition regions indicated by different colors. (**E**) Genome-wide distribution of long homopolymer and dinucleotide repeat tracts in T2T-YAO v2.0, including polyA/T (⩾10), polyC/G (⩾7), and dinucleotide repeats (⩾ 10).

Well-resolved repeat block organization was obtained for centromeric regions across all chromosomes (fig. S6A). Within higher-order repeat (HOR), a total of 536 HOR monomer types comprising 320,424 sequences were identified, with 98.61% repeats showing high sequence consistency (<5 mismatches) relative to chromosome-specific consensus sequences (data S7).

Comparison of homologous centromeres (*36*) revealed similar overall organization but substantial haplotype divergence in the size and sequence composition of each repeat block (fig. S7), consistent with previous observations of rapid centromeric evolution and expected structural variability in human centromeres (*6, 37*). Sequence divergence was predominantly associated with inactive or degenerate repeat units surrounding canonical HOR arrays, which exhibited elevated SNP density and reduced transition-to-transversion ratios (Ti/Tv) relative to genome-wide averages, as exemplified by chromosome 1 (Fig. 3C). The short arms of acrocentric chromosomes were also completely resolved. Despite being enriched in repetitive sequences shared across chromosomes, each homologous chromosome pair retained a broadly similar overall repeats organization, while still exhibiting substantial variation in the size of individual repeat blocks, resembling the structural variability observed in centromeric regions (Fig. 3D).

Telomeric regions exhibited diverse repeat organization across chromosomes in T2T-YAO v2.0. Degenerate telomeric motifs interspersed with canonical (TTAGGG)n repeats were frequently observed near subtelomeric transitions with substantial variation in repeat composition and organization across chromosomes and haplotypes (fig. S6B), similar to previous report (*38, 39*).

A total of 2,412,242 long homopolymer loci were resolved in T2T-YAO v2.0, comprising polyA/polyT (81.70%), polyC/polyG (9.20%), and poly-dinucleotide repeats (9.10%). Homopolymer distributions were highly concordant between haplotypes but showed substantial regional variation across genomic compartments, particularly in centromeric and acrocentric regions (Fig. 3E). All these results credibly characterized repetitive genomic structures, enabling high-resolution investigation of repeat organization, centromeric diversity and structurally complex genomic loci.

## Discussion

In the post-T2T era, human genomics is entering a phase in which complete diploid genomes are being generated at unprecedented scale. Ongoing population-scale genome initiatives are rapidly expanding the catalog of sequences absent from current references, many of which originate from structurally complex and highly repetitive regions that harbor substantial human diversity (*40*). As these assemblies become incorporated into reference genomes, pangenome frameworks, variant databases, and downstream biomedical analyses, accurate evaluation of their correctness becomes increasingly important to avoid propagating technical artifacts into genomic interpretation (*41, 42*). However, despite rapid advances in assembly technology, the field still lacks a consistent framework for defining and evaluating genome correctness, particularly within repetitive regions where conventional assessment approaches frequently generate ambiguous or conflicting signals. Therefore, as complete human genomes accumulate at population scale, establishing genome correctness has emerged as the next central challenge of the post-T2T era.

A key contribution of this study is the introduction of a support-based framework that approaches assembly evaluation from a fundamentally different perspective than existing methods. Most current assessment approaches infer assembly quality from read–assembly discrepancies observed within individual sequencing platforms. Although increasingly sophisticated statistical and machine-learning models have improved interpretation of these signals, their accuracy remains constrained by alignment ambiguity, sequencing biases, and platform-specific error profiles, challenges that become particularly pronounced in repetitive genomic regions. As a result, disagreement among evaluation methods often reflects uncertainty in interpreting discrepancy signals rather than differences in assembly quality itself. In contrast, support-based evaluation is not intended as another error-detection metric. Rather, it provides an experimentally grounded definition of assembly correctness based on direct sequencing evidence. By evaluating whether genomic sequences are sufficiently supported by independent and complementary datasets, the framework establishes a common criterion that can reconcile conflicting signals among existing assessment methods and localize regions requiring further investigation. Importantly, because support evaluation and support-guided refinement are both scalable to whole-genome assemblies, this framework provides a practical foundation for establishing genome correctness as complete human genomes continue to accumulate at population scale.

Our results also highlight the importance of population-relevant complete diploid genome references. Although landmark assemblies such as T2T-CHM13 and HG002 have substantially improved human genome representation (*1, 4*), current references still incompletely capture genomic diversity in East Asian populations, particularly within centromeres, segmental duplications, multicopy genes, and other rapidly evolving genomic regions. T2T-YAO v2.0 therefore serves not only as a technically refined assembly, but also as a reliable East Asian genome resource for investigating genomic variation in regions that remain underrepresented in existing references (*40, 43*). As population-scale genome initiatives continue to expand, high-quality population-specific references will become increasingly important for accurate benchmarking of variant discovery, construction of pangenome references, and development of future personalized genome resources.

Several limitations should be acknowledged. First, support-based evaluation remains dependent on read alignment and therefore inherits limitations imposed by sequence similarity and alignment ambiguity. Although SAS substantially improves the identification of unsupported regions, sufficiently long and highly identical sequences may exceed the resolving capacity of current sequencing technologies. Second, unresolved homopolymer tracts and rDNA arrays illustrate that some genomic regions remain beyond the combined accuracy and read-length limits of currently available sequencing platforms. Consequently, support-based validation should be viewed as a framework for establishing correctness within the resolving power of available experimental evidence rather than an absolute proof of sequence identity. Future improvements in sequencing accuracy, read length, and orthogonal validation technologies will further extend the scope of experimentally supported genome validation.

In summary, this study establishes a framework for defining and evaluating genome correctness using direct experimental evidence and demonstrates its application to the generation of a support-validated diploid human genome. As complete human genomes accumulate across increasingly diverse populations, rigorous standards for establishing genome correctness will become essential for constructing reliable references, benchmarking genomic analyses, and enabling precision medicine (*44, 45*).

## Acknowledgments

Induced pluripotent stem cells (iPSCs) used for the T2T-YAO assembly were obtained from our Lab (*2*). The collection of original cell used to establish iPSCs was approved by the Ethical Review Committee of Linfen Central Hospital, China (Approval No. 2022-20-1). The collection and storage of human samples were registered with and approved by the Human Genetic Resources Administration of China (HGRAC). Written informed consents were obtained from the participants. During the preparation of this work, the author(s) used Deepseek and chatGPT in order to improve language fluency, clarity, and readability. After using this tool or service, the author(s) reviewed and edited the content as needed and take(s) full responsibility for the content of the publication.

## Author contributions

YK conceived and designed the project. ZH implemented the software SAS. YC conducted the correction of T2T-YAO. SG, CS, performed sample preparation and sequencing. YC, ZH, CS, and SG contributed equally to this work. YY, YL, XY, YT, JC, RL, JW, SEA and YH participated in data analyses and discussions regarding the presentation of the data, and YK, ZG, JH, SEA and JY wrote the manuscript. All the authors agree on the manuscript.

## Data availability

The raw sequencing data for T2T-YAO v2.0 generated in this study have been deposited in the GSA for Human database at the China National Center for Bioinformation (CNCB) under accession number HRA011075, publicly accessible at https://ngdc.cncb.ac.cn/gsa-human. The raw data for v1.1 and both parents, released on August 16, 2023, are available from the same URL under accession number HRA004987. The T2T-YAO v1.1 and v2.0 genome assemblies are available in the Genome Warehouse at CNCB (https://ngdc.cncb.ac.cn/gwh) under the following accession numbers: v1.1: GWHDQZJ00000000 (maternal), GWHDOOG00000000 (paternal); v2.0: GWHGEYC00000000.1 (maternal), GWHGEYB00000000.1 (paternal). All genome sequences and associated datasets can be freely downloaded without application and are also publicly accessible at https://github.com/KANGYUlab/T2T-YAO-resources.

Public HG002 datasets used in this study, including ONT ultra-long reads, PacBio HiFi reads, Element sequencing reads, CenSat annotations and the HG002 v1.1 genome assembly, were obtained from the Human Pangenome Reference Consortium and associated public repositories.

The T2T-YAO datasets and public HG002 datasets used in this study are available from the URLs listed in data S8.

## Code availability

The SAS pipeline used for assembly assessment in this study is available at https://github.com/KANGYUlab/sas-pipeline. The platform-integrated window consensus genome polishing pipeline, referred to as PWC and genome-polish-pipeline in this study, is available at https://github.com/KANGYUlab/genome-polish-pipeline. Custom scripts used for haplotype-specific read processing, read filtering, polishing and assembly evaluation are provided in these repositories. Software versions and key parameters are described in the Supplementary Methods.

## Supplementary Materials

Materials and Methods

Figs. S1 to S7

Tables S1 to S3

References (*46*–*60*)

Data S1 to S8

## References and Notes

1. S. Nurk et al., The complete sequence of a human genome. Science 376, 44–53 (2022).

2. Y. He et al., T2T-YAO: A Telomere-to-telomere Assembled Diploid Reference Genome for Han Chinese. Genomics Proteomics Bioinformatics 21, 1085–1100 (2023).

3. C. Yang et al., The complete and fully-phased diploid genome of a male Han Chinese. Cell Res. 33, 745–761 (2023).

4. N. F. Hansen et al., A complete diploid human genome benchmark for personalized genomics. bioRxiv, 2025.09.21.677443 [Preprint] (2025). 10.1101/2025.09.21.677443.

5. P. Sarashetti et al., A Complete Telomere-to-Telomere Diploid Reference Genome for South Asian Population. bioRxiv, 2025.07.12.664550 [Preprint] (2025). 10.1101/2025.07.12.664550.

6. N. Altemose et al., Complete genomic and epigenetic maps of human centromeres. Science 376, eabl4178 (2022).

7. M. R. Vollger et al., Segmental duplications and their variation in a complete human genome. Science 376, eabj6965 (2022).

8. S. J. Hoyt et al., From telomere to telomere: The transcriptional and epigenetic state of human repeat elements. Science 376, eabk3112 (2022).

9. H. A. Tanudisastro, I. W. Deveson, H. Dashnow, D. G. MacArthur, Sequencing and characterizing short tandem repeats in the human genome. Nat Rev Genet 25, 460–475 (2024).

10. X. Liao et al., Repetitive DNA sequence detection and its role in the human genome. Commun Biol 6, 954 (2023).

11. H. A. Lawson, Y. Liang, T. Wang, Transposable elements in mammalian chromatin organization. Nat Rev Genet 24, 712–723 (2023).

12. F. Rodriguez-Algarra, D. M. Evans, V. K. Rakyan, Ribosomal DNA copy number variation associates with hematological profiles and renal function in the UK Biobank. Cell Genom 4, 100562 (2024).

13. A. Levchenko, A. Kanapin, A. Samsonova, R. R. Gainetdinov, Human Accelerated Regions and Other Human-Specific Sequence Variations in the Context of Evolution and Their Relevance for Brain Development. Genome Biol. Evol. 10, 166–188 (2018).

14. X. Wang et al., The paradox of extremely fast evolution driven by genetic drift in multi-copy gene systems. elife, (2025).

15. W. W. Liao et al., A draft human pangenome reference. Nature 617, 312–324 (2023).

16. Y. Gao et al., A pangenome reference of 36 Chinese populations. Nature 619, 112–121 (2023).

17. Y. Wang et al., The 1000 Chinese Pangenome empowers medical and population genetics. Nature 654, 121–130 (2026).

18. S. Majidian, D. P. Agustinho, C. S. Chin, F. J. Sedlazeck, M. Mahmoud, Genomic variant benchmark: if you cannot measure it, you cannot improve it. Genome Biol. 24, 221 (2023).

19. A. Rhie, B. P. Walenz, S. Koren, A. M. Phillippy, Merqury: reference-free quality, completeness, and phasing assessment for genome assemblies. Genome Biol. 21, 245 (2020).

20. Y. Ni, X. Liu, Z. M. Simeneh, M. Yang, R. Li, Benchmarking of Nanopore R10.4 and R9.4.1 flow cells in single-cell whole-genome amplification and whole-genome shotgun sequencing. Comput Struct Biotechnol J 21, 2352–2364 (2023).

21. A. S. Deshpande et al., Identifying synergistic high-order 3D chromatin conformations from genome-scale nanopore concatemer sequencing. Nat. Biotechnol. 40, 1488–1499 (2022).

22. S. Arslan et al., Sequencing by avidity enables high accuracy with low reagent consumption. Nat. Biotechnol. 42, 132–138 (2024).

23. A. M. Mc Cartney et al., Chasing perfection: validation and polishing strategies for telomere-to-telomere genome assemblies. Nat Methods 19, 687–695 (2022).

24. R. Poplin et al., A universal SNP and small-indel variant caller using deep neural networks. Nat. Biotechnol. 36, 983–987 (2018).

25. K. Shafin et al., Haplotype-aware variant calling with PEPPER-Margin-DeepVariant enables high accuracy in nanopore long-reads. Nat Methods 18, 1322–1332 (2021).

26. R. Poplin et al., Scaling accurate genetic variant discovery to tens of thousands of samples. bioRxiv, 201178 [Preprint] (2018). 10.1101/201178.

27. M. Mastoras et al., Highly accurate assembly polishing with DeepPolisher. Genome Res, (2025).

28. M. Smolka et al., Detection of mosaic and population-level structural variants with Sniffles2. Nat. Biotechnol. 42, 1571–1580 (2024).

29. M. R. Vollger et al., Long-read sequence and assembly of segmental duplications. Nat Methods 16, 88–94 (2019).

30. R. Lemmers et al., Cis D4Z4 repeat duplications associated with facioscapulohumeral muscular dystrophy type 2. Hum. Mol. Genet. 27, 3488–3497 (2018).

31. M. G. Ross et al., Characterizing and measuring bias in sequence data. Genome Biol. 14, R51 (2013).

32. T. M. Freeman C. Genomics England Research, D. Wang, J. Harris, Genomic loci susceptible to systematic sequencing bias in clinical whole genomes. Genome Res. 30, 415–426 (2020).

33. A. Shumate, S. L. Salzberg, Liftoff: accurate mapping of gene annotations. Bioinformatics 37, 1639–1643 (2021).

34. H. Li, Protein-to-genome alignment with miniprot. Bioinformatics 39, (2023).

35. J. Morales et al., A joint NCBI and EMBL-EBI transcript set for clinical genomics and research. Nature 604, 310–315 (2022).

36. L. Corda, S. Giunta, Chromosome-specific centromeric patterns define the centeny map of the human genome. Science 389, eads3484 (2025).

37. G. A. Logsdon et al., The variation and evolution of complete human centromeres. Nature 629, 136–145 (2024).

38. M. Lycka et al., TeloBase: a community-curated database of telomere sequences across the tree of life. Nucleic Acids Res. 52, D311–D321 (2024).

39. Z. Stephens, J. P. Kocher, Characterization of telomere variant repeats using long reads enables allele-specific telomere length estimation. BMC Bioinformatics 25, 194 (2024).

40. H. Jeong et al., Structural polymorphism and diversity of human segmental duplications. Nat. Genet. 57, 390–401 (2025).

41. N. Tekkey et al., Quantifying reference alignment bias in functional genomics analyses. Cell Rep Methods, 101461 (2026).

42. M. J. Lin, S. Iyer, N. C. Chen, B. Langmead, Measuring, visualizing, and diagnosing reference bias with biastools. Genome Biol. 25, 101 (2024).

43. T. Lou et al., Complete and haplotype-resolved maps of genomic and epigenetic discordance in monozygotic twins. bioRxiv, 2025.10.24.684490 [Preprint] (2025). 10.1101/2025.10.24.684490.

44. M. J. Khoury, J. Evans, W. Burke, A reality check for personalized medicine. Nature 464, 680–680 (2010).

45. D. M. Nyaga, R. E. Zaied, O. K. Silander, M. A. Black, J. M. O’Sullivan, Beyond single references: pangenome graphs and the future of genomic medicine. Front Genet 16, 1679660 (2025).

46. S. Koren et al., Canu: Scalable and accurate long-read assembly via adaptive κ-mer weighting and repeat separation. Genome Res. 27, gr.215087.215116 (2017).

47. C. Jain, A. Rhie, N. F. Hansen, S. Koren, A. M. Phillippy, Long-read mapping to repetitive reference sequences using Winnowmap2. Nat Methods 19, 705–710 (2022).

48. M. Vasimuddin, S. Misra, H. Li, S. Aluru, in 2019 IEEE International Parallel and Distributed Processing Symposium (IPDPS). (2019), pp. 314–324.

49. H. Cheng, M. Asri, J. Lucas, S. Koren, H. Li, Scalable telomere-to-telomere assembly for diploid and polyploid genomes with double graph. Nat Methods 21, 967–970 (2024).

50. M. Kolmogorov, J. Yuan, Y. Lin, P. A. Pevzner, Assembly of long, error-prone reads using repeat graphs. Nat. Biotechnol. 37, 540–546 (2019).

51. J. T. Robinson, H. Thorvaldsdottir, D. Turner, J. P. Mesirov, igv.js: an embeddable JavaScript implementation of the Integrative Genomics Viewer (IGV). Bioinformatics 39, btac830 (2023).

52. K. H. Chao et al., Combining DNA and protein alignments to improve genome annotation with LiftOn. Genome Res. 35, 311–325 (2025).

53. A. Shumate, S. Salzberg, LiftoffTools: a toolkit for comparing gene annotations mapped between genome assemblies [version 2; peer review: 2 approved]. F1000Research 11, (2024).

54. M. Tarailo-Graovac, N. Chen, Using RepeatMasker to identify repetitive elements in genomic sequences. Curr Protoc Bioinformatics Chapter 4, 4 10 11–14 10 14 (2009).

55. G. Formenti et al., Gfastats: conversion, evaluation and manipulation of genome sequences using assembly graphs. Bioinformatics 38, 4214–4216 (2022).

56. G. Benson, Tandem repeats finder: a program to analyze DNA sequences. Nucleic Acids Res. 27, 573–580 (1999).

57. H. Iseric, C. Alkan, F. Hach, I. Numanagic, Fast characterization of segmental duplication structure in multiple genome assemblies. Algorithms Mol. Biol. 17, 4 (2022).

58. I. Numanagic et al., Fast characterization of segmental duplications in genome assemblies. Bioinformatics 34, i706–i714 (2018).

59. H. Li, Minimap2: pairwise alignment for nucleotide sequences. Bioinformatics 34, 3094–3100 (2018).

60. K. J. Karczewski et al., The mutational constraint spectrum quantified from variation in 141,456 humans. Nature 581, 434–443 (2020).

