## Supplementary Materials for "Approaching an Error-Free Diploid Human Genome Using a Support-Based Validation Framework"

**The PDF file includes:**

Materials and Methods
Figs. S1 to S7
Tables S1 to S3
References

**Other Supplementary Materials for this manuscript include the following:**

Data S1 to S8

### Materials and Methods

Sample and sequencing

iPS cell line culture

Induced pluripotent stem cells (iPSCs) used for the T2T-YAO assembly were cultured in mTeSR™ PLUS medium (StemCell Technologies, Catalog No. 05825) in vitronectin (1:100 dilution, GIBCO, Catalog No. A14700) coated 6-well plates passaged at a 1:10 ratio using 0.02% EDTA (Versene, GIBCO, Catalog No. 15040-066).

ONT ultra-long library preparation and sequencing

ONT ultra-long DNA were extracted using the Grandomics BAC-long DNA Kit. The Blue Pippin system (Sage Science, Beverly, MA) was used to retrieve large DNA fragments by gel cutting. Approximately 8–10 μg of genomic DNA was selected (> 50 kb) with the SageHLS HMW library system (Sage Science), and then processed using the Ligation sequencing 1D Kit (Catalog No. SQK-LSK114, Oxford Nanopore Technologies, Oxford, UK) according the manufacturer's instructions. DNA libraries (approximately 400 ng) were constructed and sequenced on the PromethION (Oxford Nanopore Technologies) at the Genome Center of Grandomics (Wuhan, China). ONT sequencing data was generated in the pod5 format and underwent quality control using dorado in sup mode V5.0.0. This process filtered out low-quality fail reads (reads with a quality score lower than 7) to generate high-quality pass reads. These pass reads could be directly used for subsequent assembly.

Pacbio Revio® SPRQ library preparation and sequencing

High molecular weight genomic DNA was prepared by the CTAB method and followed by purification with Grandomics Genomic kit. According to the method description provided in the SMRTbell® prep kit 3.0 kit manual, the SPRQ libraries were prepared for sequencing. Sequencing data was subjected to quality control using SMRT Link V25.1 with default parameters.

Element AVITI library preparation and sequencing

For Element AVITI sequencing, PCR-free libraries were generated using the Elevate Enzymatic Library Prep Kit (Cat. No. 830-00009) and Long UDI Adapter Kit Set A (Cat. No. 830-00010), following the manufacturer's protocols. Libraries were pooled, denatured, and sequenced using the AVITI 2×150 Sequencing Kit CB UltraQ (Cat. No. 860-00018) on the AVITI platform.

Pore-C library preparation and sequencing

Cells are fixed with formaldehyde (1%) to preserve 3D chromatin structure, quenched with glycine, and washed. Crosslinked chromatin is digested with DpnII (New England Biolabs, R0543L), adjacent DNA fragments are ligated by T4 DNA ligase (Vazyme, N103-01). After reverse crosslinking, DNA is purified and prepared using the ONT Ligation Kit (SQK-LSK114) for end-repair and adapter ligation. Libraries are loaded on a PromethION at the Genome Center of Grandomics (Wuhan, China).

Reads mapping

Haplotype-specific long reads phasing and mapping

Before mapping, The ONT (>30kb) and SPRQ (>3k) reads were phased using splitHaplotype (https://github.com/KANGYUlab/genome-polish-pipeline, modified from the 'splitHaplotigs' script in Canu(*46*) with command: splitHaplotype -fastq -R long.fastq.gz -H mat.hapmer.meryl haplotype-Mat.fastq -H pat.hapmer.meryl haplotype-Pat.fastq -A unknown.fastq. Haplotype-specific reads were aligned to the corresponding haplotype assembly sequence using winnowmap(*47*) with parameters '-k 15 -W asm.meryl.repetitive_k15.txt -ax map-pb/map-ont -Y'. Haplotype- unassigned reads were aligned to diploid assembly sequences.

NGS reads mapping

Element reads were aligned to diploid assembly sequences using BWA-MEM(*48*).

SAS assessment

Preprocessing of multi-platform sequencing data

To ensure the fidelity of all downstream analyses, we implemented a rigorous, technology-specific preprocessing and filtering pipeline for all input alignment (BAM) files.

For short-read data, alignments were subjected to stringent selection criteria, retaining only those reads that exhibited a perfect match to the reference genome (specified by the SAM tag NM:i:0) and a complete absence of soft or hard clipping events. This foundational step was critical for establishing a high-confidence baseline, essential for the subsequent high-precision identification of base-level errors.

For long-read data generated on PacBio HiFi (Revio) and Oxford Nanopore (ONT) platforms, our analyses were exclusively restricted to primary alignments. Secondary and supplementary alignments were deliberately excluded to mitigate analytical ambiguities arising from reads mapping to multiple genomic loci. Furthermore, to manage artifacts associated with read fragmentation while preserving genuine structural information, we designed and applied an adaptive filter to regulate split alignments. The maximum permissible number of splits for a given read was calculated dynamically based on its length, following the equation:

$$\text{Max\_splits} = 3.0 + (\text{Read\_length\_kb} \times 0.05)$$

This strategy effectively removes reads with excessive fragmentation, which often indicates low-quality sequencing or chimeric artifacts, while appropriately accommodating the natural correlation between read length and the incidence of split alignments in authentic long reads.

According to the aforementioned algorithm, read alignments were filtered with command "filter.sh long-read/short-reads -i in.bam -o filter.bam". All scrip file is available at https://github.com/KANGYUlab/genome-polish-pipeline). All filtered bam files from the same sequencing platform were merged.

Structural Error (SE) Detection

Framework for Structural Error Detection

Structural error (SE) detection was performed exclusively using ultra-long ONT sequencing data, leveraging its superior capacity to span large and complex genomic structures that are challenging for shorter reads. Our analytical framework integrates three independent modules, each targeting distinct structural features to provide comprehensive error identification.

By default, the assembly was scanned using non-overlapping 2 kb windows. A window was flagged as an SE if:

1. INDELs and Clipping: >40% of the local reads contained insertions or deletions ⩾50 bp detected via CIGAR string analysis, or Presence of hard or soft clipping in aligned reads.
2. Abnormal Depth: Read depth <20% or >200% of the genome-wide mean (assessed using Mosdepth).
3. Lack of Spanning Reads: No spanning reads were observed within 15 consecutive 2 kb windows (i.e., a 30 kb region).

Thresholds can be relaxed near chromosome ends (<10 kb from termini), where a window is flagged only if no or fewer than 10% of local reads spanned the region and lacked indels.

Insertion and deletion detection

Large insertions and deletions (INDEL, ⩾50 bp) were systematically identified through CIGAR string analysis within 2-kb windows across the genome. For each window, we quantified both the total count of insertion and deletion events and the number of unique reads supporting each event type. To ensure accurate representation of large deletions that may span multiple analysis windows, reads contributing deletion signals were allocated to all affected windows, thereby preventing underestimation of deletion events.

Anomalous depth profiling

Potential assembly collapses and structural misjoins were detected through systematic read depth analysis using mosdepth with 2-kb resolution windows. Genomic regions were flagged as potential structural errors if their local coverage deviated significantly from the genome-wide mean depth: regions with depth below 20% of the mean were classified as potential duplications or misjoins, while regions exceeding 200% of the mean depth were identified as potential collapses.

Clipping event analysis

Potential assembly misjoins were identified through comprehensive analysis of soft and hard clipping events within 2-kb sliding windows. A critical feature of this module is the deliberate exclusion of clipping events occurring at the precise chromosomal termini (i.e., the first and last base positions). This design choice specifically mitigates false positives arising from incomplete telomere extension in reference assemblies, thereby distinguishing genuine internal structural errors from assembly artifacts at chromosome ends.

Non-split Read Coverage (NScov%) Calculation

The Non-split Read Coverage (NScov%) was calculated to measure the genome-wide coverage provided by contiguous, non-chimeric reads. First, nanopore sequencing alignments (BAM file) were filtered with Samtools to retain only primary alignments, discarding all secondary and supplementary records. Next, any primary alignment containing an SA tag—indicative of a split-read or chimeric event—was also removed. The per-base depth of this filtered set was then calculated using mosdepth. NScov% was defined as the proportion of the reference genome covered at a depth of at least 1× by these non-split reads.

Base-level Error Detection

Framework for base-level error detection

Element reads were aligned to the diploid assembly using BWA-MEM(*48*). Alignments were filtered to retain only perfect matches (CIGAR NM:i:0) without clipping. SPRQ reads were phased and aligned similarly to ONT reads.

Base-level errors were evaluated in sliding 50 bp windows. Windows were flagged as BEs if the exact sequence was supported by reads at <10% of the mean sequencing depth. Windows with sufficient (>10% mean depth) perfect-match Element reads were excluded. Remaining regions were further validated using SPRQ and ONT data. In low-coverage regions, thresholds were relaxed to 10% of local depth under the following conditions:

1. Within 10 kb of chromosomal termini (i.e., telomeric regions);
2. Regions with degenerative GA/CT repeats (⩾90% GA/CT and ⩾60% GC content);
3. GC-rich regions (⩾90% GC content).

<u>Merqury-Based Assembly Evaluation</u>

Merqury(*19*) and Meryl(*19*) were used to assess k-mer-based quality metrics (QV), completeness, and phasing accuracy using 21-mers and 31-mers. 21-mers and 31-mer were counted in the child, maternal and paternal read sets, and haplotype-specific mers (hapmer) were created using Merqury scripts with the command 'hapmers.sh mat.meryl pat.meryl son.meryl'. The evaluation followed the methods described by McCartney et al5(*23*) The binary k-mer set comprised Element and SPRQ k-mers set with >1 occurrence, while the trinary k-mer set additionally included ONT read k-mers with >4 occurrences.

<u>Variant Calling (SVs and SNVs)</u>

Structural variants-like errors were called using Sniffles2(*28*) (ONT), Flagger(*15*) (ONT), and NucFlag (https://github.com/logsdon-lab/NucFlag) (SPRQ), with default parameters. Sniffiles2 SVs with supportive reads less than 60% were removed. Base-level SNV-like errors were identified using DeepVariant(*24*) (ONT, --model_type=ONT_R104; Element, --model_type=WGS), Deeppolisher(*27*) (SPRQ), and GATK HaplotypeCaller(*26*) (Element). Variants with genotype quality (GQ) <20 were excluded. Heterozygous calls were also filtered out except for haplotype-aware calls from Deeppolisher.

<u>Error Correction</u>

<u>Structural error correction</u>

SE windows were first corrected using variant calls from Sniffles2 when available. In cases without confident calls, ONT reads spanning the SE and 5kb-10kb flanks were reassembled using Hifiasm(*49*) (--ont) or Flye(*50*) (--nano-raw). If required, flanking regions were extended to 50–100 kb for resolving large collapses or duplications. Reassembled contig was aligned back to reference, and the corrected sequences were substituted accordingly. Manual assessment and correction with Integrative Genomics Viewer(*51*) (v2.6) is often necessary for SE of clipping or complex errors.

Telomeric region were detected using Teloscope with default parameters [https://doi.org/10.1093/bioinformatics/btac460]. For telomere extension, ONT reads spanning the telomere boundary and extending ⩾20 kb inward from the subtelomeric side were reassembled using Hifiasm (--ont) or Flye (--nano-raw). If required, spanning regions were extended to 50–100 kb for resolving large collapses or duplications. Reassembled contigs were aligned back to reference for telomere extension.

In telomeric regions with misalignment or heterogeneity, ONT reads were classified into two haplotypic groups based on heterozygous SNP patterns within target region. Specifically, reads spanning target region were extracted from aligned BAM files, and variants—including SNPs and indels—were identified by parsing alignment CIGAR strings. High-confidence heterozygous SNPs with balanced allele frequencies (⩾0.2) were selected as informative markers. For each read, a haplotype vector was constructed by encoding the alleles at these heterozygous sites. Reads were then classified via hierarchical clustering based on the Hamming distances between haplotype vectors. Finally, reads with sparse coverage at informative sites were assigned to the resulting clusters based on their average sequence similarity to the classified read. Major-allele ONT reads were reassembled for telomeres extension.

Corrections were followed by haplotype-aware re-alignment and re-evaluation with SAS. The process was repeated until no SEs were detected outside rDNA clusters.

Base-level error polishing (PWC pipeline)

BE windows were classified into homopolymer-associated (h-BE) and non-homopolymer-associated (nh-BE).

Candidate error sites were evaluated for reference sequence support across sequencing platforms. Sites were excluded as false positives if any platform showed >55% of supporting reads (⩾10 reads) supporting the reference sequence, with an additional window read-depth cutoff of ⩽300 for the Element platform.

**nh-BEs** were corrected using consensus sequences from supporting reads in the order of platform priority: Element > SPRQ > ONT. Consensus sequences were constructed using samtools consensus with adaptive thresholds based on sequence composition. For regions with AG or CT frequency <90%, normal mode was applied with consensus threshold 0.70. For regions with AG or CT frequency ⩾90%, relaxed mode was used with consensus threshold 0.50. Consensus sequences were accepted only if minimum coverage (⩾10 reads) and consensus threshold requirements were met. Where consensus could not be reached from Element, SPRQ and ONT platforms were sequentially attempted. When all platforms failed, local ONT reassembly was performed for nh-BE regions.

**h-BEs** were polished using Element or SPRQ consensus only, due to ONT's known limitations in homopolymer resolution. The same consensus construction parameters and thresholds as for nh-BE were applied.

Polishing iterations continued until the number of detected BEs plateaued.

SAS assessment of HG002 assembly

All sequencing data and annotations of HG002 used in this study are publicly available. No new data were generated in this study. ONT UL reads were obtained from https://s3-us-west-2.amazonaws.com/human-pangenomics/index.html?prefix=T2T/scratch/HG002/sequencing/ont/12_1_22_R1041_ULCIR_HG002_dorado0.4.0/, HiFi reads were obtained from https://s3-us-west-2.amazonaws.com/human-pangenomics/T2T/HG002/assemblies/polishing/HG002/v1.0/mapping/hifi_revio_pbmay24/hg002v1.0.1_hifi_revio_pbmay24.bam convert to fastq, Element reads were obtained from https://s3-us-west-2.amazonaws.com/human-pangenomics/index.html?prefix=T2T/scratch/HG002/sequencing/element/trio/HG002/). CenSat region was obtained from https://s3-us-west-2.amazonaws.com/human-pangenomics/T2T/HG002/assemblies/annotation/centromere/h g002v1.1_v2.0/hg002v1.1.cenSatv2.0.noheader.bb. The HG002 v1.1 were obtained from https://s3-us-west-2.amazonaws.com/human-pangenomics/T2T/HG002/assemblies/hg002v1.1.fasta.gz.

According to the part of "Reads mapping", "SAS assessment" and "Variant Calling (SVs and SNVs)" described earlier in the Methods, sequencing reads of HG002 sample from three platforms were aligned to HG002v1.1, followed by BAM filtering, SAS evaluation and Variant Calling (Sniffles2 for SVs calling, Deeppolisher for SNVs calling).

Benchmarking with simulated SVs and SNVs

For structural errors (SEs), we randomly introduced 4,900 large indels (52–9,998 bp) and 50 inversions (1,067–4,852 bp) into the hg002v1.1 and YAO v2.0 assemblies using SURVIVOR (v1.0.7), excluding acrocentric chromosomes. For base-level errors (BEs), we introduced 18,000 SNPs and 2,000 small indels into both assemblies using SimuG (v1.0.1) with parameters -snp_count 18000 -indel_count 2000 -ins_del_ratio 0.5. Haplotype-specific long reads (ONT and HiFi), together with element reads, were aligned to the spike-in genomes according to the previous parameter process. We applied SAS for both SE and BE detection, and used Sniffles2 and DeepPolisher as comparison methods for SE and BE detection, respectively. We used BEDTools intersect to obtain the counts of true positives (TP), false negatives (FN), and false positives (FP), and subsequently calculated recall, precision, and the F1 score. The coordinates of residual error regions were lifted over from the original assemblies to the spike-in genomes using minimap2 (v2.27) and transanno (v0.4.4). For evaluation of simulated error detection, to avoid potential bias caused by positional shifts between the simulated SNV coordinates and the actual mismatch or misjoin sites based on reads alignment, we expanded the Deeppolisher, SAS-SE and SAS-BE regions by one detection window on both flanks (50 bp for BE detection and 2 kb for SE detection). All downstream performance assessments were conducted based on these expanded regions.

Identification of identical regions (IRs): IRs were identified using genmap (K=150, E=0) to generate a genome-wide k-mer mapping frequency bedgraph. Regions with non-unique k-mer coverage were identified, and their end positions were extended to 150 bp downstream from the start position, constrained by chromosome boundaries. Overlapping intervals were merged using bedtools merge to generate the final IR.

Annotation

Genome annotation

We generated annotations for T2T-YAO v2.0 maternal and paternal haplotypes separately by mapping genes and transcripts from the latest annotation of T2T-CHM13 (GCF_009914755.1-RS_2025_08) to T2T-YAO v2.0 each haplotype.

Before global annotation, we masked some complex genomics regions include the V(D)J gene segments and the ribosomal DNA (rDNA) arrays to minimize mapping errors in these regions. We lifted V(D)J gene annotations from the RefSeq GFF of the T2T-CHM13 (GCF_009914755.1-RS_2025_08) onto T2T-YAO v2.0 each haplotype using Liftoff(*33*), and curated manually to ensure accurate regions of V(D)J gene. Total rDNA arrays were identified by aligning reference rDNA sequence KY962518.fasta to the T2T-YAO v2.0 using nucmer (--maxmatch -l 31 -c 100; delta-filter -i 96 -l 1000).

Then, we used Liftoff (-chroms chroms.txt -copies -sc 0.95 -exclude_partial -polish) to map all features from T2T-CHM13 (GCF_009914755.1-RS_2025_08) onto T2T-YAO v2.0 each haplotype, excluding V(D)J gene segments and rDNA arrays. We then manually removed the previously identified complex genomic regions (V(D)J gene segments and rDNA arrays) from the generated annotations. This process yielded 56919 genes on MAT and 56546 genes on PAT.

To obtain comprehensive annotation of T2T-YAO v2.0, particularly for protein-coding genes, we supplemented the annotation with GRCh38 MANE (v1.4) annotations. Following the extra-copy search submodule strategy in (*52*), we aligned MANE protein sequences to T2T-YAO each haplotype using miniport(*34*) with default parameters. Comparing the miniprot gene feature to the previously annotations lifted from T2T-CHM13, we filtered out any miniprot features that overlapped ⩾10% of their own length with an existing gene feature or spanned more than two existing adjacent gene loci to avoid two genes were annotated in the same genomic location. This procedure added 15 genes to MAT and 29 to PAT.

We further refined the annotation coordinates using the protein-maximization algorithm in LiftOn, generating a comprehensive and accurate T2T-YAO v2.0 gene annotation dataset for downstream analyses. Synteny of the genes between T2T-CHM13 and T2T-YAO were plotted using LiftoffTools(*53*) synteny module (liftofftools synteny -r <reference.fa> -t <target.fa> -rg <reference.gff3> -tg <target.gff3>).

### Centromere and Telomere Annotation

Centromere regions were annotated using RepeatMasker(*54*) (based on Dfam39 library) with default parameters, and only hits with scores >50 were retained. The α-satellite arrays were identified via HumAS-HMMER(*37*) with default parameters. The sequence composition of human centromeric α-satellite HOR arrays was visualized using CenMAP (v0.5.1) (https://github.com/logsdon-lab/CenMAP). The orientation, and organization of centromere protein B (CENP-B) binding motifs were annotated using Genomic Centromere Profiling (GCP) pipeline(*36*). Telomeric repeats were detected and quantified using Teloscope with default parameters(*55*) [https://doi.org/10.1093/bioinformatics/btac460].

### Identification of Segmental Duplications (SDs)

To identify segmental duplications (SDs), we implemented the detection workflow established for the CHM13 T2T genome assembly(*7*). The full pipeline for the masking and SD identification steps followed the framework provided at https://github.com/mrvollger/assembly_workflows/ (under workflows/sedef.smk). Within this integrated workflow, common repeats were masked with RepeatMasker (v4.1.9)(www.repeatmasker.org) and Tandem Repeats Finder (v4.09)(*56*). BISER (v1.4)(*57*) was employed within the pipeline as a technical advancement over SEDEF(*58*), providing optimized and accelerated characterization of duplication structures.

### ClinVar Coordinate Liftover

Since the original ClinVar coordinates are based on the GRCh38 (GCF_000001405.40) reference, we performed a coordinate liftover to the YAO assembly using the standard transanno protocol. First, a sequence alignment "chain" between GRCh38 and YAO was established using minimap2 (v2.30-r1290-dirty)(*59*). Subsequently, the coordinate transformation was executed via transanno (v0.4.4; https://github.com/informationsea/transanno) to project the ClinVar variants onto the YAO assembly.

### SD Synteny

Following the established methodology(*7*) for defining syntenic regions between T2T-CHM13 and GRCh38, we constructed multiple genome alignments using Cactus (v2.9.9; https://github.com/ComparativeGenomicsToolkit/cactus). The resulting alignments were stored in Hierarchical Alignment Format (HAL). Syntenic blocks were then extracted and analyzed using halSynteny (v2.2; https://github.com/ComparativeGenomicsToolkit/hal) to identify orthologous relationships within complex and repetitive segmental duplication regions.

Analysis of Genomic Variations

T2T-YAO v2.0 was separated into paternal (pat) and maternal (mat) haplotypes. In addition to an internal comparison between the two haplotypes, each haplotype was independently aligned against both the T2T-CHM13v2.0 and GRCh38.p14.

For each pairwise comparison, whole-genome alignment was performed using Minimap2 (v2.28)26 (-cx asm5 --cs, --eqx). Syntenic one-to-one alignments were extracted from the resulting PAF files, sorted, and then processed with paftools.js call (-q 60 -L 1000) to generate a comprehensive VCF file. The VCF file was then filtered to retain only SNPs and short indels (1-3bp) for each chromosome comparison pair. Ti/Tv ratios, SNP and short indels density were calculated from these one-to-one aligned regions.

Functional consequences of all identified variants were annotated using the Ensembl Variant Effect Predictor (VEP, v114, https://github.com/Ensembl/ensembl-vep) with respect to the corresponding reference annotation for each pair. Putative loss-of-function (pLoF) variants were identified by the Loss-Of-Function Transcript Effect Estimator(*60*) plugin and flagged as 'High-Confidence' (HC).

Pore-C chromosomal configuration data analysis

Haplotype phasing were validated with Pore-C contact maps using wf-pore-C v1.3.0 (https://github.com/epi2me-labs/wf-pore-c) with parameters '--cutter 'DpnII' –ref asm.diploid.fasta'. Contact maps were visualized in Juicebox v2.20 (https://github.com/aidenlab/Juicebox).

A

B

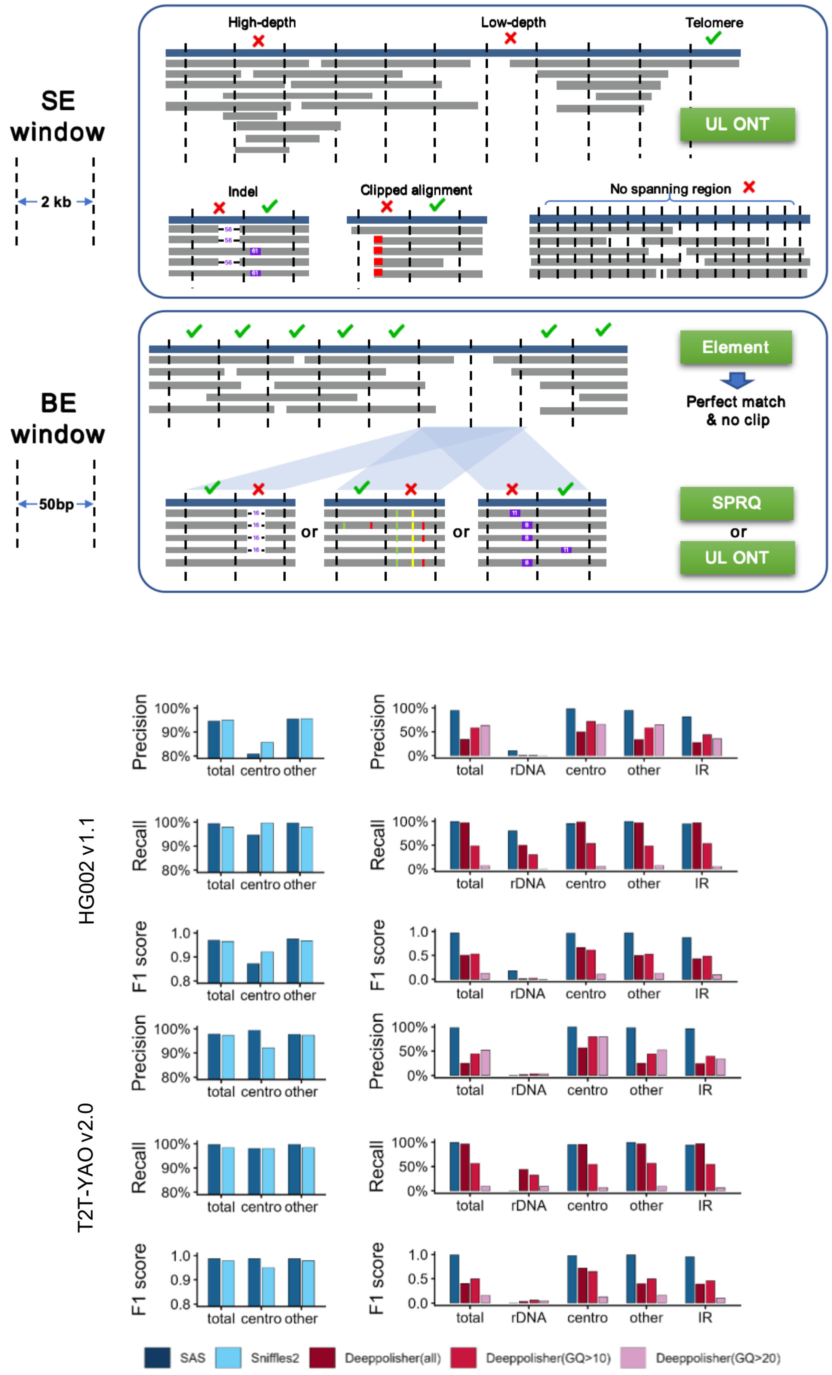

**Fig. S1. Definition and genome-wide detection of residual errors by SAS.** (**A**) SAS evaluates assembly correctness based on alignment support from sequencing reads. In structural error (SE) windows (2 kb), support is assessed using ultra-long Oxford Nanopore (ONT) reads, requiring consistent spanning alignments and normal depth; regions with clipped alignments, absence of spanning reads, or abnormal depth are flagged as structural inconsistencies. In base-level error (BE) windows (50 bp), support is evaluated using high-accuracy reads, requiring mismatch-free and unclipped alignments from at least one platform (Element, SPRQ, or ONT). (**B**) Precision–recall analysis using simulated spike-in errors demonstrates high accuracy of SAS for both SE and BE detection in HG002 and T2T-YAO v2.0.

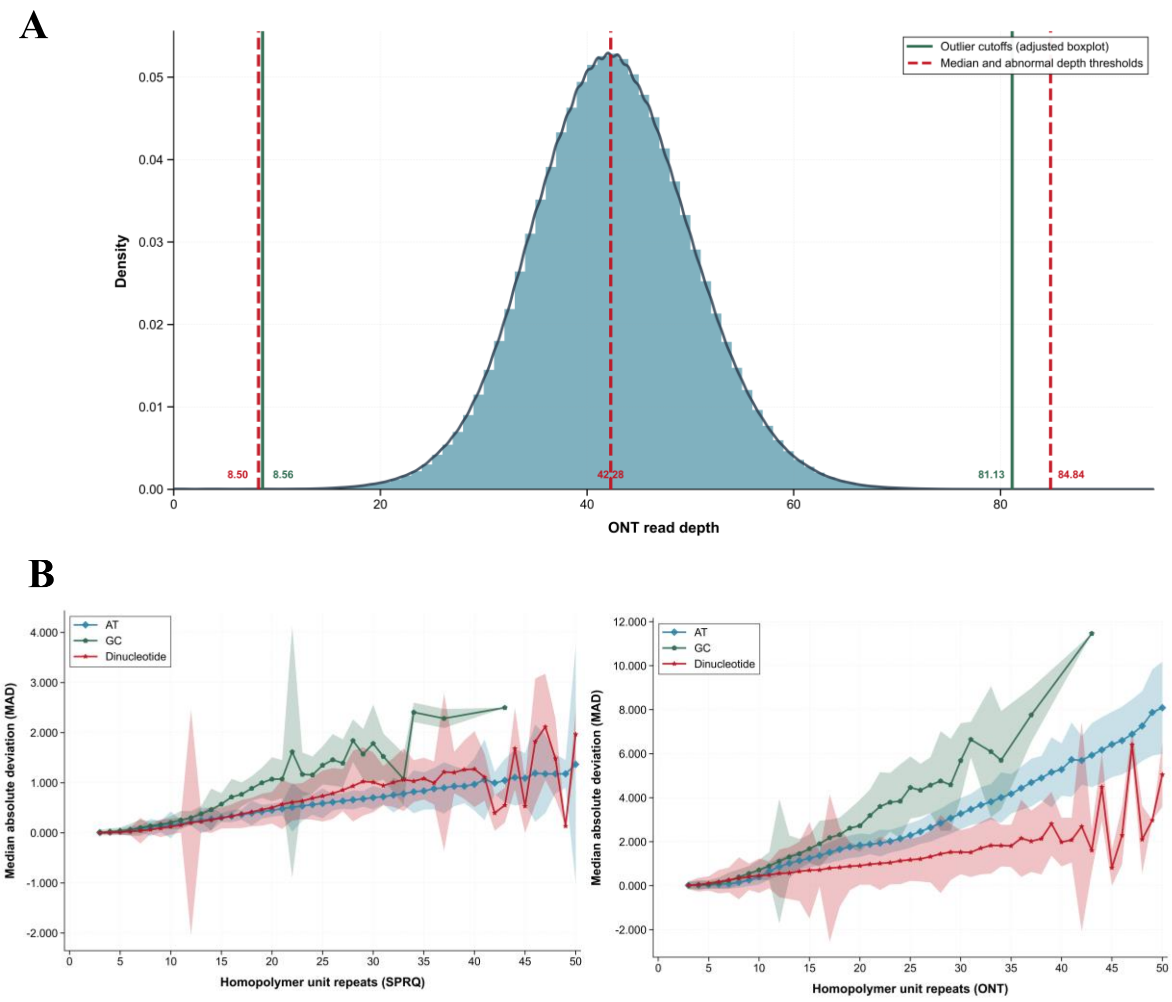

**Fig. S2. Statistics of read depth and homopolymer length deviation.** (**A**) Genome-wide distribution of ONT read depth. Green solid lines indicate the upper and lower outlier cutoffs calculated using adjusted boxplot (based on MedCouple), while red dashed lines denote the median and thresholds for abnormal depth. (**B**) Median absolute deviation (MAD) of the number of repeat units along homopolymer tracts in SPRQ (left panel) and ONT (right panel) reads.

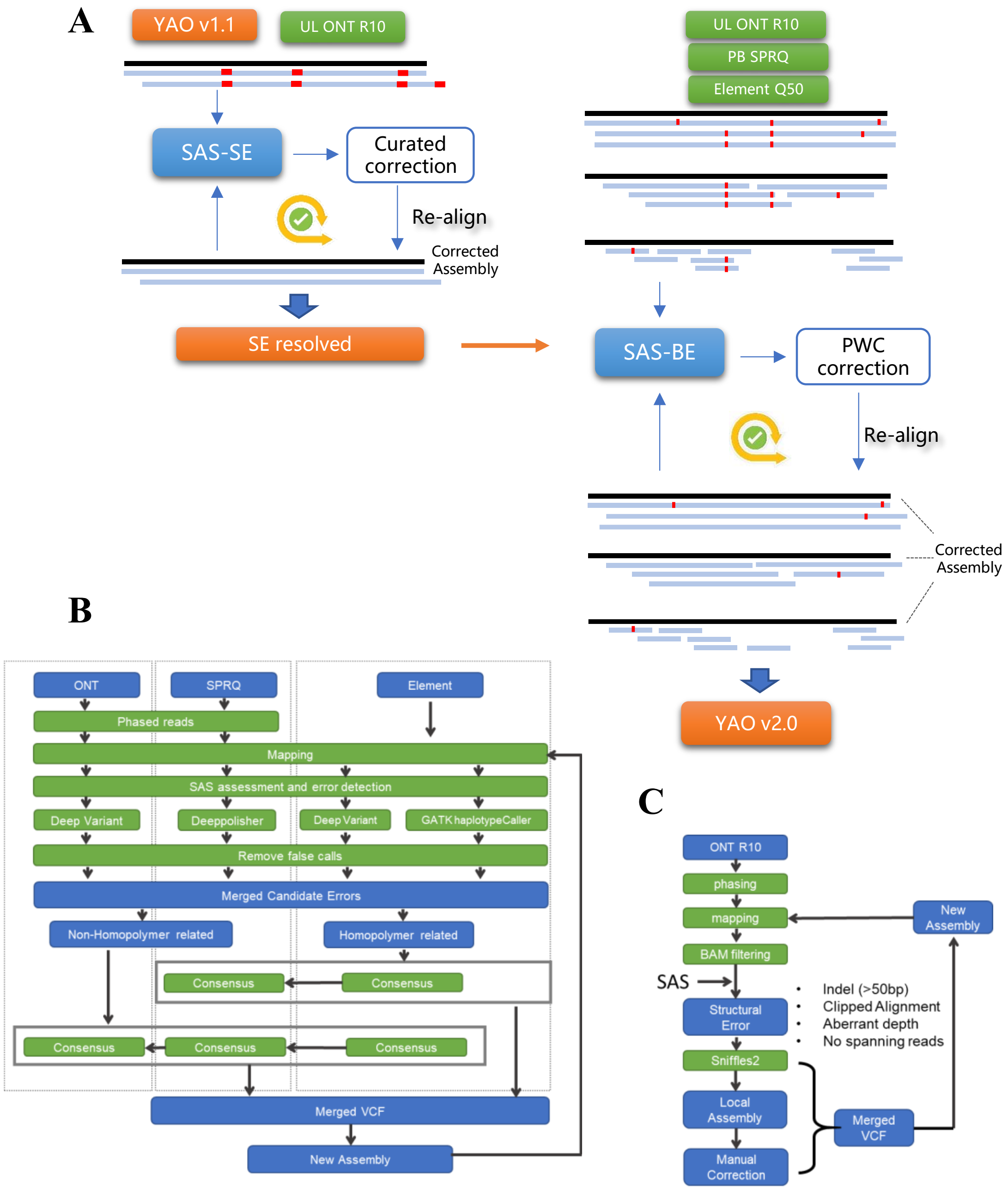

**Fig. S3. SAS-guided iterative correction of residual assembly errors.** (**A**) Residual inconsistencies in T2T-YAO v1.1 are resolved through a two-stage, SAS-guided correction framework. Structural errors (SEs) are first identified using ultra-long Oxford Nanopore (ONT) reads and corrected through targeted curation and local reassembly, followed by read realignment and re-evaluation by SAS until structural support is continuous (SE-resolved). The updated assembly is then subjected to base-level evaluation (SAS-BE) using multi-platform data (ONT, SPRQ, and Element). Base-level inconsistencies are corrected using a platform-integrated window consensus (PWC) strategy, and reads are iteratively realigned and reassessed. This sequential, structure-first refinement minimizes alignment artifacts and enables convergence toward a fully supported assembly. Wide and narrow red bar indicate SE an BE, respectively. (**B**) Schematic Overview of the Structural Error (SE) Correction Pipeline. (**C**) Schematic overview of the PWC (Platform-integrated Window Consensus) pipeline.

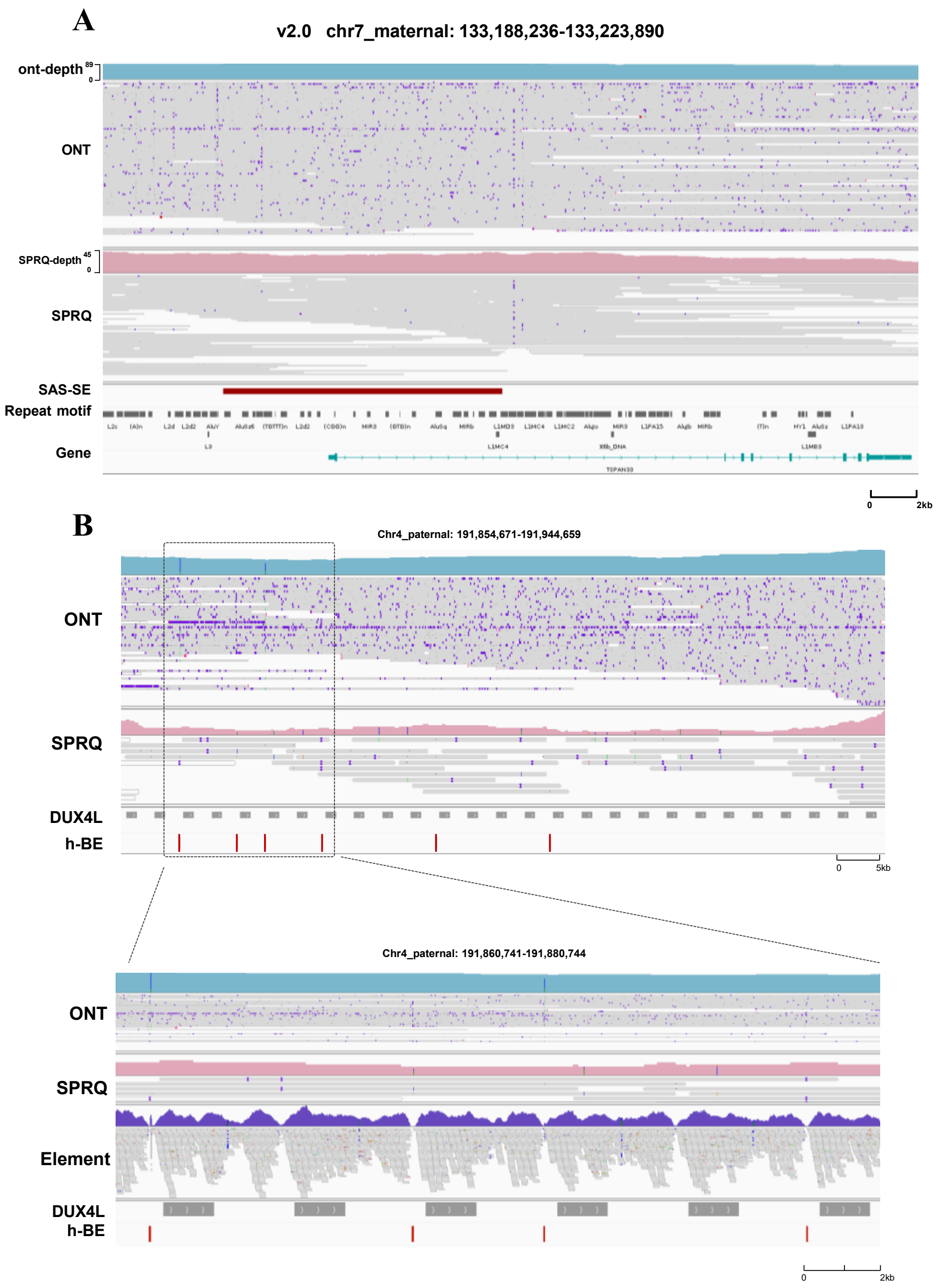

**Fig. S4. Remaining inconsistencies in T2T-YAO v2.0.** (**A**) The remaining SE (structural error) region on maternal chromosome 7 with abnormal ONT read depth. The sequencing depth (blue peak for ONT and pink peak for SPRQ) and read alignment from ONT, SPRQ platforms are displayed across the SE region in maternal chromosome 7. The SAS-SE track highlight the location of abnormal ONT read depth (dark red) detected by SAS pipeline. The bottom tracks highlight the genomic features, including repeat motif and genes in this region. (**B**) Example of unresolved homopolymer sites in the D4Z4 repetitive region on chromosome 4q35. The sequencing depth and read alignment from multiple sequencing platforms (ONT, SPRQ and Element) are displayed with peaks above each track representing the sequencing depth at each position. The gray horizontal bars indicate the sequencing reads, with colored vertical bars denoting mismatches in these reads. The bottom tracks highlight the position of DUX4L genes and unresolved h-BE.

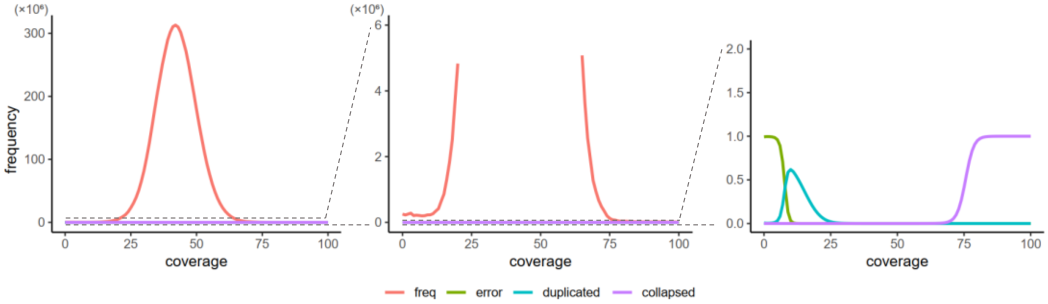

**Fig. S5. Genome-wide ONT read depth distribution in T2T-YAO v2.0.** Most regions follow a near-Gaussian distribution, whereas predicted misassemblies (collapsed or duplicated regions) are confined to extreme coverage tails, predominantly corresponding to repetitive sequences such as rDNA arrays.

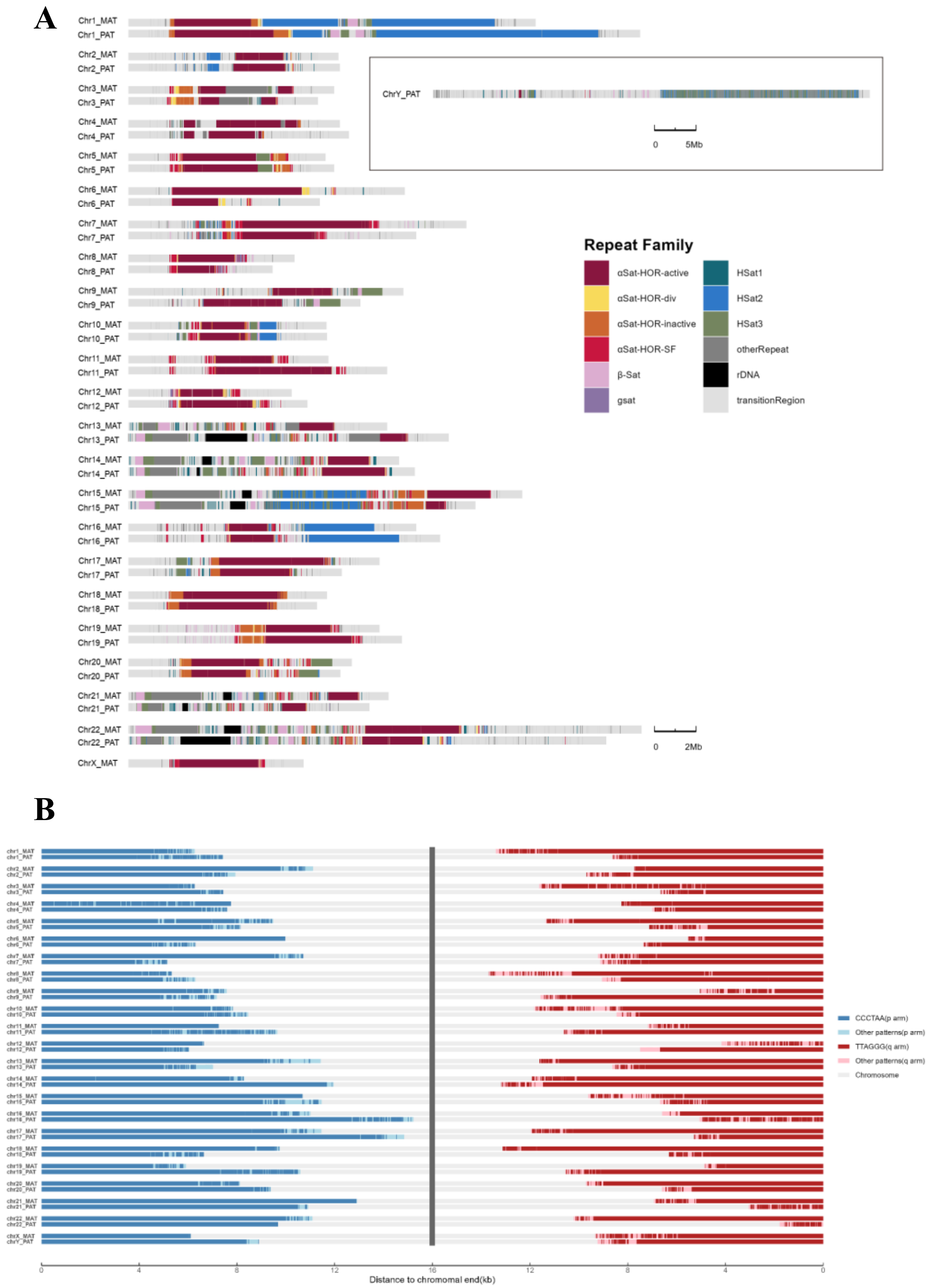

**Fig. S6. Structural characterization of centromeres and telomere in the polished T2T-YAO v2.0 assembly.** (**A**) Annotation of centromeric regions in all the 46 chromosomes. Repeat content is visualized as stacked bars by annotated repeat family. Each pair of centromeres displays a unique repeat composition and organization, with notable differences between haplotypes. The inset shows a zoom-out view of the Y chromosome, which is full of repeat sequences. (**B**) Telomeric repeat content and extension across all 46 chromosomal termini. Each bar represents the distance from the chromosomal terminus, with colored segments indicating specific 6-mer tandem repeats. Canonical (TTAGGG)n (in long arm) and complementary (CCCTAA)n (in short arm) motifs are shown in dark red and blue, along with shallow colors indicate non-canonical telomeric-like 6-mer repeats.

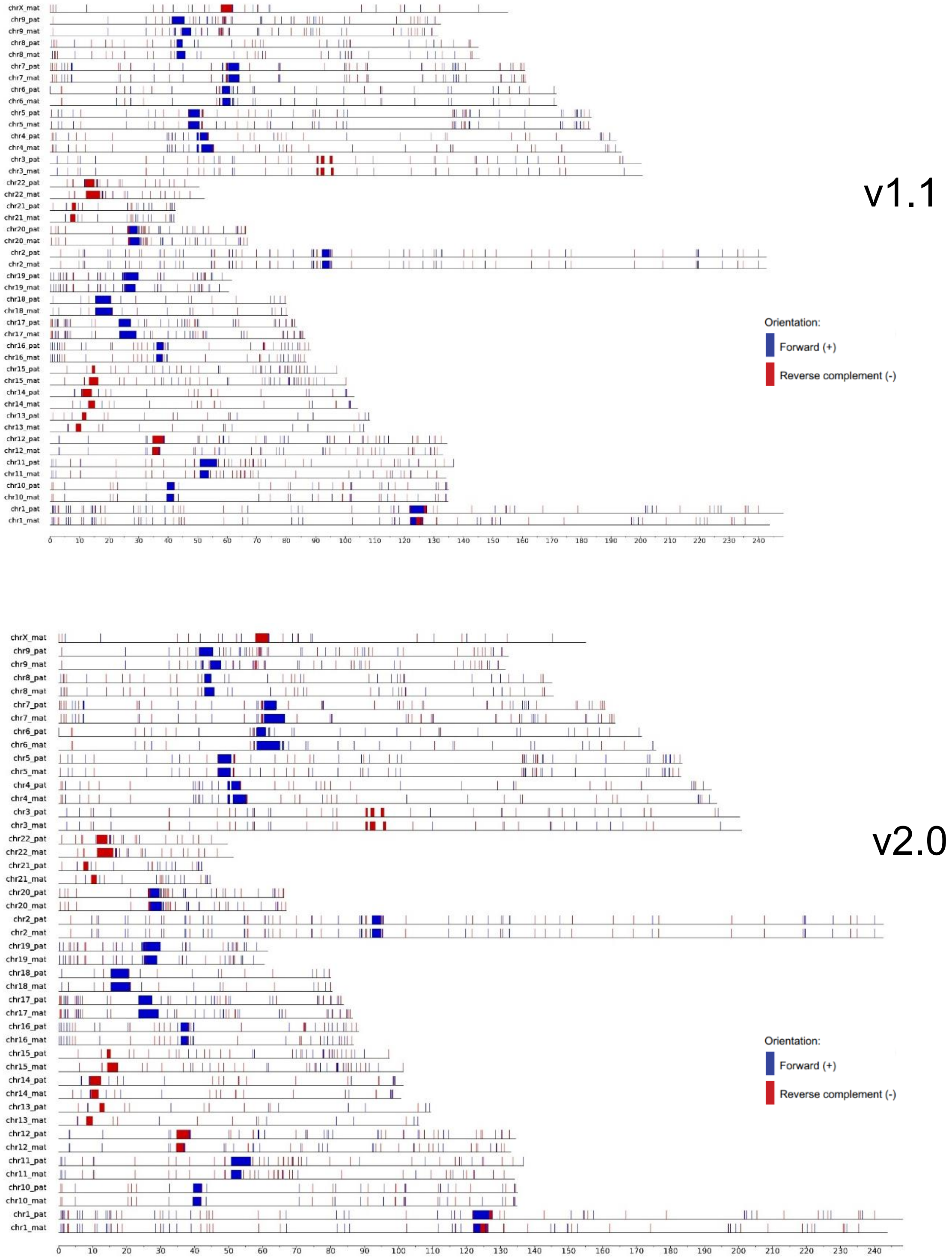

**Fig. S7. Comparison of centromere infrastructure of T2T-YAO v1.1 and v2.0 via Genomic Centromere Profiling (GCP).**

**Table S1. Sequencing datasets and phasing statistics for the evaluation of the T2T-YAO assembly**

| Platform | total base | total reads | N50 |
|---|---|---|---|
| ONT_total(>30k) | 260,204,701,389 | 2,719,918 | 123,536 |
| ONT_mat | 115,834,438,677 | 1,110,641 | 134,245 |
| ONT_pat | 110,226,498,350 | 1,059,358 | 133,913 |
| ONT_unknown | 34,143,764,362 | 549,919 | 67,361 |
| RevioSPRQ_total(>3k) | 306,486,031,653 | 15,535,705 | 19,584 |
| RevioSPRQ_mat | 80,354,712,631 | 4,034,614 | 19,773 |
| RevioSPRQ_pat | 77,454,694,722 | 3,890,390 | 19,763 |
| RevioSPRQ_unknown | 148,676,624,300 | 7,610,701 | 19,398 |
| Element | 320,980,626,300 | 2,139,870,842 | - |
| Pore-C | 172107855509 | 36,725,533 | 5,581 |

**Table S2. Comparative quality metrics of the T2T-YAO assembly assessed by Merqury**

| | | v1.1 | | | v2.0 | | |
|---|---|---|---|---|---|---|---|
| | | **mat** | **pat** | **Both** | **mat** | **pat** | **Both** |
| **Merqury-** binary | **QV-21mer** | **66.48** | **68.68** | **67.42** | **74.84** | **78.23** | **76.18** |
| | err-kmer | 14252 | 8304 | 22556 | 2085 | 921 | 3006 |
| | switch err% | 0.0107 | 0.0254 | — | 0.0017 | 0.0038 | — |
| | Completeness% | 97.77 | 93.708 | 99.839 | 97.777 | 93.71 | 99.85 |
| | **QV-31mer** | **60.97** | **62.73** | **61.74** | **67.24** | **69.04** | **68.03** |
| | err-kmer | 74864 | 48221 | 123085 | 17694 | 11262 | 28956 |
| | switch err% | 0.0219 | 0.0413 | — | 0.005 | 0.0072 | — |
| | Completeness% | 96.78 | 92.607 | 99.806 | 96.792 | 92.61 | 99.826 |
| **Merqury-** trinary | **QV-21mer** | **67.7888** | **69.854** | **68.6823** | **inf** | **inf** | **inf** |
| | err-kmer | 10543 | 6331 | 16874 | 0 | 0 | 0 |
| | switch err% | 0.0107 | 0.0254 | — | 0.0017 | 0.0038 | — |
| | Completeness% | 97.7896 | 93.7264 | 99.8638 | 97.7966 | 93.7281 | 99.875 |
| | **QV-31mer** | **62.4363** | **64.2025** | **63.2153** | **96.2941** | **88.7314** | **91.0965** |
| | err-kmer | 53376 | 34337 | 87713 | 22 | 121 | 143 |
| | switch err% | 0.0219 | 0.0413 | — | 0.005 | 0.0072 | — |
| | Completeness% | 96.81 | 92.636 | 99.841 | 96.823 | 92.639 | 99.861 |

### Table S3. Gene annotation of T2T-YAO v2.0 liftoff from T2T-CHM13

| | Gene Type | CHM13 | Unique | Success % | Extra copy | Failures | Failure % | Total | MANE Unique | MANE Extra |
|---|---|---|---|---|---|---|---|---|---|---|
| Maternal | Protein coding | 19987 | 19842 | 99.27% | 182 | 145 | 0.73% | 20024 | 11 | 4 |
| | LncRNA | 15511 | 15291 | 98.58% | 625 | 220 | 1.42% | 15916 | — | — |
| | Pseudogene | 16599 | 16197 | 97.58% | 674 | 402 | 2.42% | 16871 | — | — |
| | Others | 3899 | 3806 | 97.61% | 302 | 93 | 2.39% | 4108 | — | — |
| | Total | 55996 | 55136 | 98.46% | 1783 | 860 | 1.54% | 56919 | 56930 | 56934 |
| Paternal | Protein coding | 19238 | 19087 | 99.22% | 213 | 151 | 0.78% | 19300 | 9 | 20 |
| | LncRNA | 16072 | 15773 | 98.14% | 1258 | 299 | 1.86% | 17031 | — | — |
| | Pseudogene | 16059 | 15651 | 97.46% | 649 | 408 | 2.54% | 16300 | — | — |
| | Others | 3733 | 3675 | 98.45% | 240 | 58 | 1.55% | 3915 | — | — |
| | Total | 55102 | 54186 | 98.34% | 2360 | 916 | 1.66% | 56546 | 56555 | 56575 |
| Diploid | Protein coding | 20090 | 19975 | 99.43% | — | 115 | 0.57% | 39324 | 14 | 30 |
| | LncRNA | 16388 | 16116 | 98.34% | — | 272 | 1.66% | 32947 | — | — |
| | Pseudogene | 16993 | 16654 | 98.01% | — | 339 | 1.99% | 33171 | — | — |
| | Others | 3906 | 3855 | 98.69% | — | 51 | 1.31% | 8023 | — | — |
| | Total | 57377 | 56600 | 98.65% | — | 777 | 1.35% | 113465 | 113479 | 113509 |

**Data S1.**

Supplementary Figures. Genomic distribution of assembly inconsistencies across individual chromosomes before and after polishing.

**Data S2 to S8**

Data S2. Structural Errors Identified by Multiple Tools Before and After Correction
Data S3. Benchmark of SAS in detecting assembly errors.
Data S4. Chromosome-Wise Assembly Quality Metrics Before and After Polishing
Data S5. GC and GA/CT Content of Remaining Erroneous 31-mers in T2T-YAO v2.0
Data S6. The copy number and location of each gene in T2T-CHM13, maternal and paternal haplotype of the T2T-YAO v2.0
Data S7. Specific HOR units in T2T-YAO v2.0
Data S8. Public datasets and assemblies used in this study
